# Computational design of *de novo* integrated domains enables rational control of pathogen effector recognition in plant NLR immune receptors

**DOI:** 10.64898/2026.07.10.737686

**Authors:** Y. Xi, A.H. Bucknell, J.L. Watson, A. Maqbool, J.W. Bennett, I. Goreshnik, D. Vafeados, M. Garcia Sanchez, G. Knight, R. Zdrzałek, C.A. Rodney, I. Saado, C.E. Stone, E.K. Turley, D.S. Yu, A. Gentle, L.S. Ryder, X. Yan, V. Were, J. Heddle, D. Baker, P.M.F. Emmrich, N.J. Talbot, M.J. Banfield, A.R. Bentham

## Abstract

The rapid evolution of plant pathogens poses a persistent threat to global agricultural sustainability, often outpacing the discovery and deployment of natural disease resistance genes. While bioengineering of plant intracellular immune receptors (NLRs) offers a potential solution, developing bespoke immune recognition remains constrained by the laborious characterisation of natural receptors and plant-pathogen interactions. Here, we describe a programmable framework that leverages generative AI protein design tools, RFdiffusion and ProteinMPNN, to design de novo integrated domains (IDs) against diverse pathogen effectors. By integrating these bespoke binders into the modular rice blast Pik-1/Pik-2 NLR receptor chassis, we successfully engineer recognition of a non-cognate virulence factor (effector) from the Panama disease pathogen, *Fusarium oxysporum* f. sp. *cubense* Tropical Race 4. Functional assays in *Nicotiana benthamiana* demonstrate that these de novo domains facilitate specific effector perception and initiate immune signalling, while structural and biophysical analyses confirm that de novo integrated domains maintain high structural fidelity to the initial designs and associate with their targets via the predicted interaction interfaces. Additionally, our findings provide orthogonal evidence for the role of integrated domains in regulation of NLR signalling, demonstrating integration of de novo IDs can either trigger autoactivity or, in some cases, lead to effector-mediated repression of cell death. By decoupling immune perception from natural evolutionary history through deploying AI-designed sensory domains, this work establishes a design-lead framework for generation of programmable plant immune receptors, providing a new avenue for bioengineering crops against emerging pathogens.

## Introduction

The relentless evolution of plant pathogens represents a critical challenge for maintaining global food security, as traditional disease resistance genes are frequently overcome by rapid diversification of pathogen populations^1,2^. While bioengineering the plant immune system has emerged as a promising solution to this threat, the field remains hindered by a significant technical hurdle: the development of bespoke resistance is currently a slow and labour-intensive process, requiring extensive characterisation of host-pathogen interactions, and is often difficult to translate between different pathosystems^1,3,4^. Recent advances in computational protein design have expanded the scope of protein engineering, enabling creation of de novo proteins with desirable properties^5–9^. Studies using protein design pipelines utilising tools for backbone design and inverse folding, such as RFdiffusion and ProteinMPNN, have demonstrated the ability to generate high-affinity protein binders against a range of complex targets, including the SARS-CoV-2 spike protein^7^, the insulin receptor^8^, and snake venoms^9^, often achieving high affinity interactions without the need for extensive experimental maturation^7^. Despite their transformative potential, these tools have not been applied to crop bioengineering^10,11^.

The bioengineering of plant immunity often relies on the manipulation of nucleotide-binding leucine-rich repeat (NLR) receptors^1,12,13^. NLRs are intracellular immune receptors that recognise pathogen virulence proteins (effectors) secreted during host invasion, activating effector-triggered immunity (ETI), which often culminates in programmed cell death to isolate the pathogen and prevent disease^14,15^. A particularly compelling chassis for bioengineering effector recognition is the rice Pik paired NLR complex (Pik-1/Pik-2). Multiple allelic variants of the Pik pair exist in rice, such as Pikp-1/Pikp-2 and Pikm-1/Pikm-2, that have evolved to recognise alleles of AVR-Pik^16,17^, an effector from the blast fungus *Magnaporthe oryzae* belonging to the *Magnaporthe* AVRs and ToxB-like (MAX) effector family^18^. Recognition of AVR-Pik by the Pik pair occurs via direct interaction of the effector with an integrated heavy metal-associated (HMA) domain embedded in the Pik-1 receptor, with polymorphisms in the HMA domain determining the recognition specificity of different Pik-1 alleles for specific AVR-Pik variants^16,19,20^. Effector recognition by Pik-1 in the presence of Pik-2 results in hypersensitive cell death and subsequent disease resistance^16,19,21^. Architecturally, the Pik-1 receptor is distinctive among NLRs carrying integrated domains (IDs); while most IDs are appended to the termini of the receptor, the Pik-1^HMA^ domain is positioned between the N-terminal coiled-coil (CC) and the central NB-ARC domains^16,17,22^. Crucially, this HMA domain is solely responsible for both effector perception and subsequent activation of the Pik-1/Pik-2 immune complex^3,23,24^, rendering it both necessary and sufficient for defense response activation. This functional autonomy facilitates a ‘plug-and-play’ modularity, where the effector recognition profile of Pik-1 can be manipulated by interchanging this domain^3,25–27^. Previous efforts have successfully expanded the recognition profile of the Pik-1 receptor by grafting VHH nanobodies^3,25^ (such as anti-GFP or anti-mCherry VHH) into the Pik-1 framework or by incorporating host HMA-containing proteins, such as OsHIPP43 and OsHIPP19, to confer recognition of non-cognate MAX effectors, such as Pwl2^27^, or unrecognised alleles of AVR-Pik, such as AVR-PikC and AVR-PikF^26^. Collectively, these studies have highlighted the exceptional modularity of this system, opening new avenues for incorporating novel protein domains to alter pathogen recognition. However, to date the Pik receptors have not been modified to enable recognition of non-MAX effectors or effectors from phytopathogens other than *M. oryzae*. Thus, whether Pik receptors can serve as a versatile scaffold for the recognition of structurally distinct effectors from diverse phytopathogens remains an open question.

In this study, we utilised an established AI-driven protein design framework comprising RFdiffusion, ProteinMPNN and AlphaFold2, to generate protein binders against three effectors, one MAX effector and two non-MAX effectors, and integrated these binders into the Pik-1 chassis where they would function as bespoke, new-to-nature integrated domains (Figure 1 A). FoSSP17, is a small, secreted non-MAX effector from *Fusarium oxysporum* f. sp. *cubense* Tropical Race 4 (*Foc* TR4), causal agent of Panama disease in banana^28^. This effector plays an important role in virulence of *Foc* TR4 by suppressing pattern-triggered immunity, reactive oxygen species accumulation, and callose deposition^28^, but no cognate resistance gene has been identified. Notably, FoSSP17 localises to both the nucleus and plasma membrane in plants^28^, highlighting its suitability for NLR-based resistance engineering. Previous field trials of transgenic Cavendish lines expressing the NBS-LRR gene *RGA2* have demonstrated that NLR-based resistance to *Foc* TR4 is achievable in principle^29,30^, however bioengineering of NLRs for *Foc* TR4 effector recognition has not been reported. Furthermore, FoSSP17 homologs are broadly conserved across *Fusarium* species and extend into related members of the *Hypocreomycetidae*^28^, raising the possibility that engineered recognition of this effector could confer resistance with utility beyond a single Panama disease pathotype^11^, making FoSSP17 a particularly desirable target for engineering.

**Figure 1.**
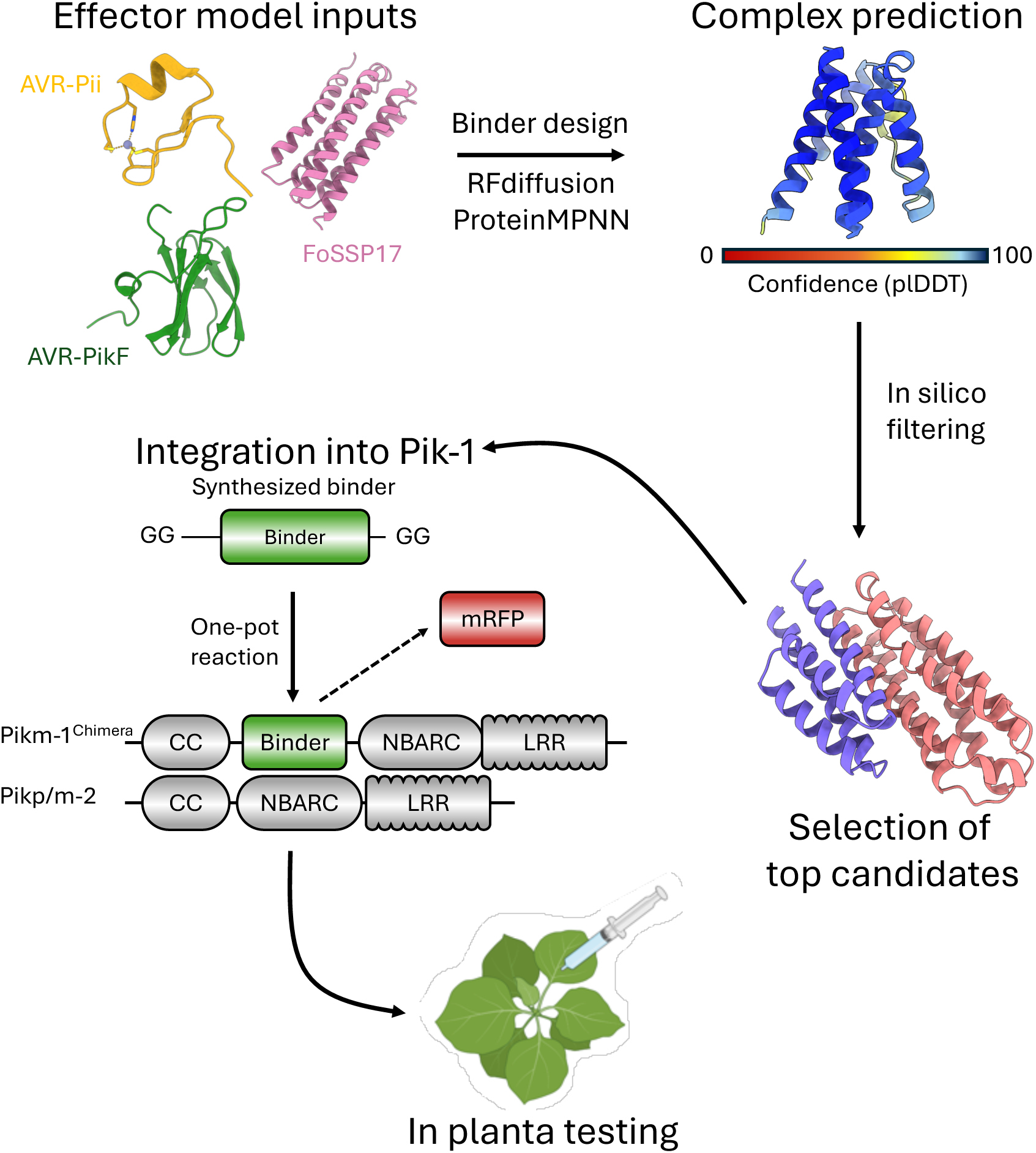
Workflow from computational design of de novo integrated domains to implementation in the Pikm-1 NLR. Structural information of the AVR-PikF, AVR-Pii and FoSSP17 effectors are used as inputs for binder generation with RFdiffusion. ProteinMPNN is subsequently used for inverse folding before structure prediction with AlphaFold2. In silico filtering using AlphaFold2 confidence metrics (pLDDT, ipTM and ipAE) performed to generate a pool of top candidate sequences for integration in the Pikm-1 receptor. Using a tailored Golden Gate (GG) Pikm-1 integrated domain acceptor vector, binders are synthesized with complementary GG overhangs and cloned into the native Pik-1 integrated domain site to generate Pikm-1 binder chimeras for in planta testing.

Through biochemical characterisation and in planta assays, we demonstrate that de novo binder integration into the Pik-1 chassis can confer FoSSP17-dependent cell death responses in *N. benthamiana*. Biophysical and structural characterisation of binder/effector complexes shows that these designed domains bind their target with high fidelity to the original design, regardless of the resulting cell death phenotype, indicating that binding and downstream transduction of the binding event into receptor activation represent two separable steps in NLR engineering. Furthermore, observation of cell death repression phenotypes within our panel of binders provides orthogonal evidence that the integrated domain site functions as a sensitive regulatory hub, in which effector binding can either trigger or suppress NLR activation depending on the specific binder used. By decoupling immune perception from natural evolutionary history using AI-designed integrated domains, this work establishes a design-first framework for generation of programmable plant immune receptors and demonstrates the utility of de novo protein design as both an effector recognition engineering strategy and a tool for probing the determinants of NLR activation.

## Results

### De novo protein binders can be generated against pathogen effectors

Our aim was to develop novel sensors to bind pathogen virulence proteins that could function as IDs when incorporated into the Pik-1 chassis. We selected two effectors from *M. oryzae*, AVR-PikF and AVR-Pii, and FoSSP17 from *Foc* TR4. Our rationale was to include a MAX effector with previously demonstrated engineered resistance (AVR-PikF), a non-MAX effector from *M. oryzae* (AVR-Pii), and an effector from an economically significant pathogen outside of *M. oryzae*, *Foc* TR4 (FoSSP17). As outlined in Figure 1, we performed protein binder design campaigns with RFdiffusion^7^ using existing structural data for AVR-PikF^31^ and AVR-Pii^32^ as input targets; for FoSSP17 we used a high-confidence structure prediction generated with AlphaFold2^33,34^ v1.5.5 as implemented in ColabFold (plDDT > 0.90; pTM >0.85), because no experimental structural data existed for this effector at the time (Figure S1). For AVR-PikF and FoSSP17, a large-scale RFdiffusion binder design campaign was refined by partial diffusion and screened by yeast surface display to identify binding candidates (See methods). For AVR-Pii 256 backbones were generated with RFdiffusion as implemented in ColabDesign^35^, which were used as templates for inverse folding with ProteinMPNN^6^ that provided eight sequences per backbone, resulting in a total of 2048 binder sequences. Binder and effector sequences were subjected to structure prediction with AlphaFold2 and filtered based on confidence metrics (plDDT >0.85; ipTM >0.80; ipAE < 5.0) to obtain final designs for experimental validation in planta.

### De novo designed immune sensors trigger cell death in response to pathogen effectors through the Pik-1 NLR chassis

We utilised well-established cell death assays in *N. benthamiana* to determine possible phenotypes arising from inserting these putative de novo IDs into Pik-1^3,16,24^. To do this, we substituted the Pikm-1^HMA^ domain with binder sequences (Figure 1) and tested the Pikm-1 chimeras in *N. benthamiana* alongside either the Pikm-2 or Pikp-2 helper, with the Pikp-2 NLR having previously been demonstrated to be more tolerant of different domain integrations into Pik-1^3,20^. Chimeric Pikm-1 NLRs were co-expressed with their Pik-2 helpers in *N. benthamiana* via *Agrobacterium*-mediated infiltration in the presence and absence of their target effector and leaves were imaged five days post infiltration. For testing the AVR-Pii binders, Pikm-1 and Pikm-2 co-expressed with AVR-PikD were used as positive controls for cell death; for testing the AVR-PikF and FoSSP17 binders, Pikm-1_OsHIPP43 co-expressed with Pikp-2 and Pwl2 was used as a positive control due to its robust cell death response in *N. benthamiana*^27^.

Of the 44 binders generated against AVR-PikF and integrated in Pikm-1, none were observed to yield any cell death in an effector-dependent manner (Figure S2). The majority of phenotypes observed either showed no recognition of AVR-PikF (*n* = 14), or autoactive cell death upon integration of the binders into Pikm-1 in the absence of AVR-PikF (*n* = 30) (Figure S2 A and C). For the five binders designed against AVR-Pii tested, we also observed phenotypes corresponding to autoactive cell death (*n* = 1) and no recognition (*n* = 2). We observed indications of effector-dependent cell death responses (*n* = 2) for some binders when integrated into Pikm-1 and co-expressed with AVR-Pii and Pikm-2 (Figure S2 B and C). However, these results were inconsistent across biological replicates and therefore we are unable to confidently attribute cell death to effector recognition. A total of 46 FoSSP17 binders were integrated into Pikm-1 (herein referred to as Fobinders) for testing in *N. benthamiana*. Of the 46 Pik-1 chimeras tested, we observed recognition events for two Fobinders, Fobinder1 and Fobinder4, which triggered cell death in an FoSSP17-dependent manner when co-expressed with the Pikp-2 helper (Figure 2 A). Similarly, we observed enhanced cell death for Fobinder2 and Fobinder3 in the presence of FoSSP17, indicative of effector recognition, although these constructs showed evidence of autoactivation in the absence of the effector (Figure 2 A and C). The remaining 42 Fobinder constructs either resulted in autoactivation of cell death in the absence of the effector (*n* = 31) or failed to elicit cell death in the presence of FoSSP17 (*n* = 11; Figure S2 C). Western blot analyses demonstrated protein accumulation of 4xMyc-tagged FoSSP17, FLAG-tagged Pikm-1_Fobinder chimeras, and HA-tagged Pikp-2 (Figure S3 A and E). Taken together, these data demonstrate de novo designed IDs can facilitate effector-dependent cell death upon incorporation into Pikm-1 NLR.

**Figure 2.**
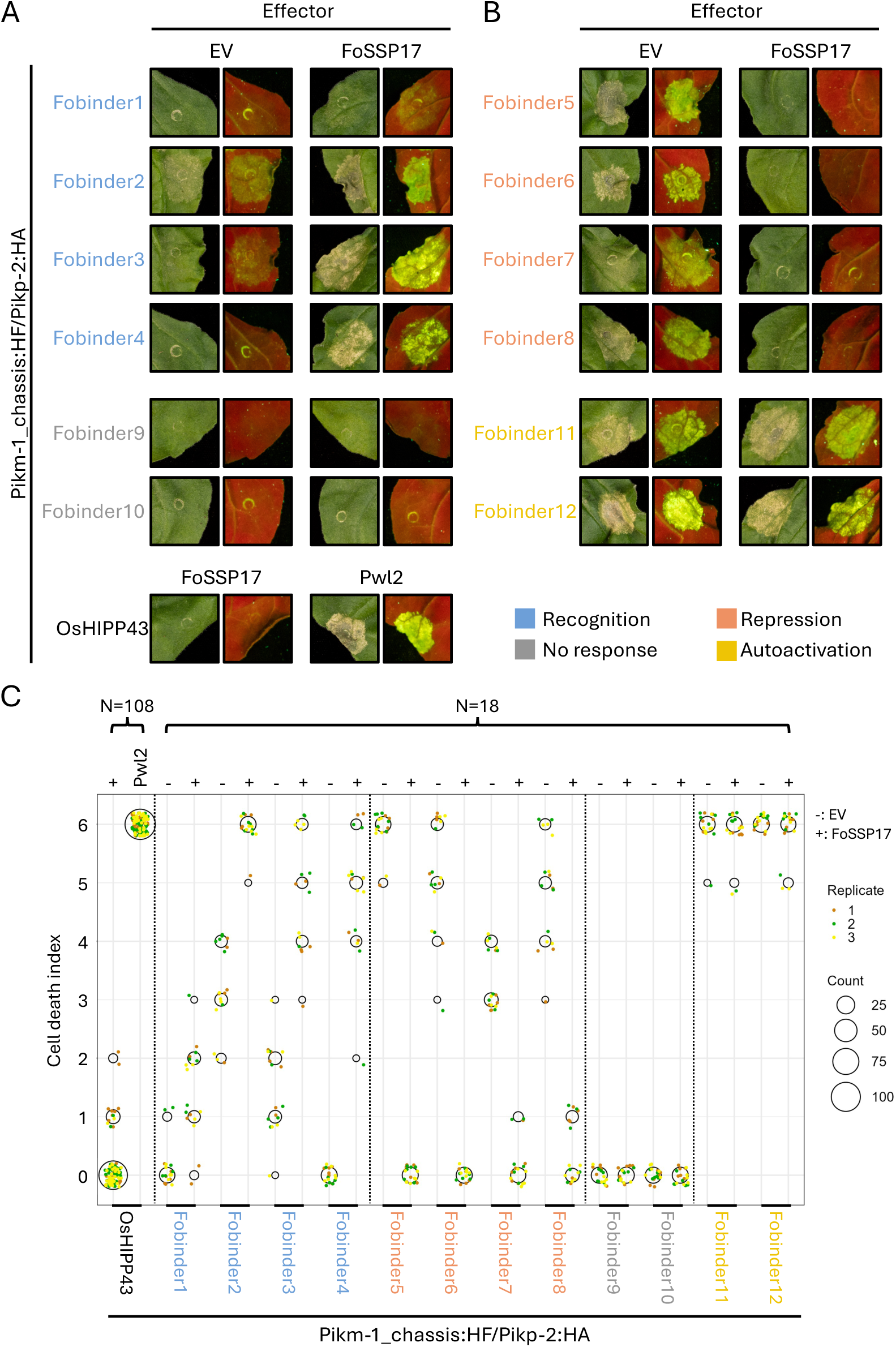
Cell death assays of Pikm-1_Fobinder chimeras in the presence and absence of FoSSP17 display binder-specific phenotypes. (A) Representative leaf images (left: white light; right: UV light) of Pikm-1_chassis:HF/Pikp-2:HA chimeras carrying Fobinder1, Fobinder2, Fobinder3 and Fobinder4 (Recognition, blue) or Fobinder9 and Fobinder10 (No response, grey), co-expressed with either empty vector (EV) or FoSSP17 and imaged five days post-infiltration. Pikm-1_OsHIPP43/Pikp-2 co-expressed with FoSSP17 or Pwl2 are shown as negative and positive controls for cell death, respectively. (B) As in (A), for chimeras carrying Fobinder5, Fobinder6, Fobinder7 and Fobinder8 (Repression, salmon) or Fobinder11 and Fobinder12 (Autoactivation, yellow). The colour key shown applies to the Fobinder labels in (A), (B) and (C). (C) Dot plot of cell death scores for all twelve Pik_Fobinder chimeras and the Pikm-1_OsHIPP43/Pikp-2 control, co-expressed with empty vector (“-”) or FoSSP17 (“+”) as indicated, or with Pwl2 for the OsHIPP43 positive control. Cell death phenotypes were scored from 0 to 6 as described previously (De la Concepcion et al., 2018). Three biological replicates (shown in different colours) were performed; the total number of replicates (N) for each group is indicated above the plot. The size of the central circle at each score is proportional to the number of replicates with that score.

### Binder integration-induced autoactivity of the Pik pair can be repressed by effector co-expression

As previously reported, the integration of different protein domains into the Pik-1 chassis can result in constitutive cell death in the presence of the Pik-2 helper NLR, without an effector being present (autoactivation)^3,26,27^. Consistent with this, the majority of binders we tested in the Pik-1 scaffold resulted in autoactivation (Figure S2 C). Interestingly, we observed several instances where autoactive cell death phenotypes could be repressed upon addition of the effector (Figure 2B; Figure 3). Fobinders 5–8, targeting FoSSP17 were demonstrated to be autoactive in the presence of Pikp-2, while co-expression with FoSSP17 in *N. benthamiana* resulted in attenuation of the autoactive cell death response (Figure 2 B and C; Figure 3 A). These data suggest the effector may interact with the binder domain in such a way as to prevent the activation of the Pik-1/Pik-2 complex. To confirm this was a FoSSP17-specific effect, Fobinders 5–8 were co-expressed with two unrelated effectors, AVR-PikD and Pwl2. Neither reproduced the repression phenotype, confirming that attenuation of cell death is FoSSP17-dependent (Figure 3 A and C). Consistent with this interpretation, an autoactive Pik pair Pi1-5C/Pi1-6C^20^ continued to cause cell death when co-expressed with FoSSP17 (Figure 3 B and C), further demonstrating that the repression is associated with integration of the designed binder and not through a general suppression of immune signalling by the effector itself.

**Figure 3.**
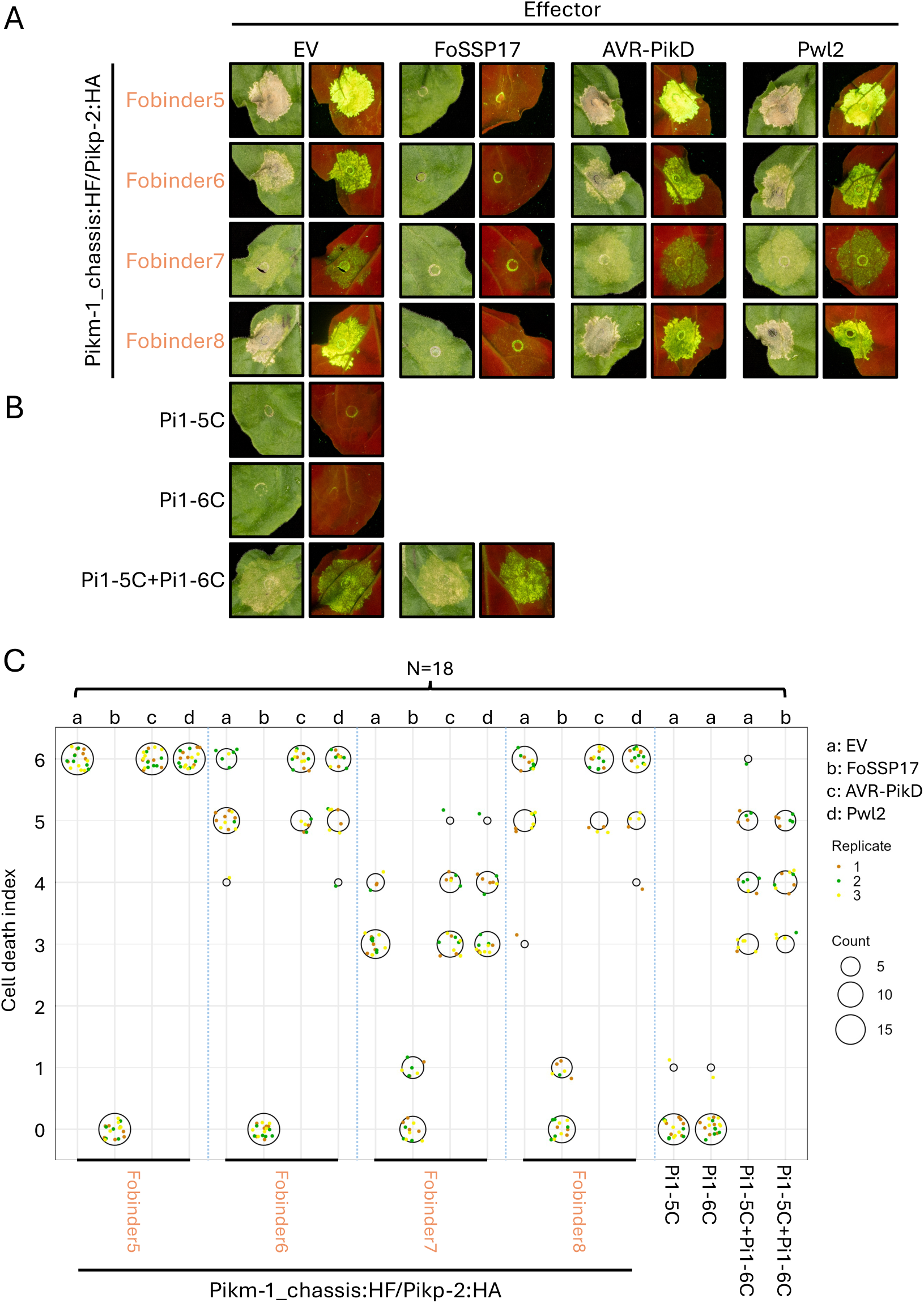
Autoactivation caused by integration of Fobinders 5-8 into Pikm-1 is specifically repressed by FoSSP17. (A) Representative leaf images (left: white light; right: UV light) of Pikm-1_Fobinder5, Fobinder6, Fobinder7 and Fobinder8 chimeras (co-expressed with Pikp-2) following co-expression with empty vector (EV), FoSSP17, AVR-PikD, or Pwl2. Cell death in the EV condition reflects the autoactivity of each Fobinder chimera; attenuation of autoactivity is observed specifically in the presence of FoSSP17, but not AVR-PikD or Pwl2. (B) Representative leaf images of the autoactive Pik pair Pi1-5C and Pi1-6C, which lacks an integrated domain, expressed alone or together, with EV or FoSSP17. Pi1-5C+Pi1-6C-triggered cell death is not repressed by FoSSP17, indicating that the repression observed for Fobinders 5-8 (A) depends on the integrated domain rather than a general suppressive effect of FoSSP17 on Pik-mediated signalling. (C) Dot plot of cell death scores for Fobinder5-8 chimeras under each effector condition (“a”: EV, “b”: FoSSP17, “c”: AVR-PikD, “d”: Pwl2), and for the Pi1-5C, Pi1-6C and Pi1-5C+Pi1-6C controls under EV (“a”) or FoSSP17 (“b”). Cell death phenotypes were scored from 0 to 6 as described previously (De la Concepcion *et al.*, 2018). Three biological replicates (shown in different colours) were performed; N is indicated above the plot. The size of the central circle at each score is proportional to the number of replicates with that score.

### In planta co-immunoprecipitation assays demonstrate that pathogen effectors associate with chimeric NLRs through their de novo domains

Given the observation of FoSSP17-dependent cell death (Figure 2) and FoSSP17-dependent repression of cell death (Figure 3) from the integration of Fobinders into Pik-1, we sought to confirm these phenotypes were a product of association of the effectors with Pik-1 chimeras through their de novo integrated domains in planta. From our cell death screening, we included the four Fobinders that resulted in effector-dependent cell death (Fobinders 1 – 4), four Fobinders that resulted in repression of autoactivity (Fobinders 5–8), two Fobinders that showed no response to FoSSP17 (Fobinder9 and 10) and two Fobinders that triggered autoactivity unable to be repressed by FoSSP17 (Fobinder11 and 12). Co-immunoprecipitation assays were performed with FLAG-tagged Pikm-1 chimeras and cognate Myc-tagged effectors. We included previously reported chimeric Pikm-1_OsHIPP43:HF co-expressed with Pwl2:Myc as a positive control^27^ and with Myc:FoSSP17 as a negative control. Constructs were expressed in *N. benthamiana* via Agrobacterium-mediated transformation with tissue harvested after 40 hours. We observed specific association between all the twelve Pikm-1_Fobinder chimeras with FoSSP17 (Figure 4), with no association with the Pwl2 effector (Figure S4), demonstrating that de novo designed integrated domains are facilitating association of FoSSP17 with Pikm-1 chimeras. These data provide evidence that FoSSP17-dependent cell death, and indeed FoSSP17-dependent repression, results from association of the effector with the Fobinder-Pik-1 chimeras. However, the association of FoSSP17 with all Fobinders tested, including those that demonstrated no effector-dependent cell death responses, suggests that phenotypic differences in cell death between constructs reflect each binder’s capacity to translate recognition into activation of the NLR upon interaction with FoSSP17, rather than being determined solely by its ability to associate with the effector.

**Figure 4:**
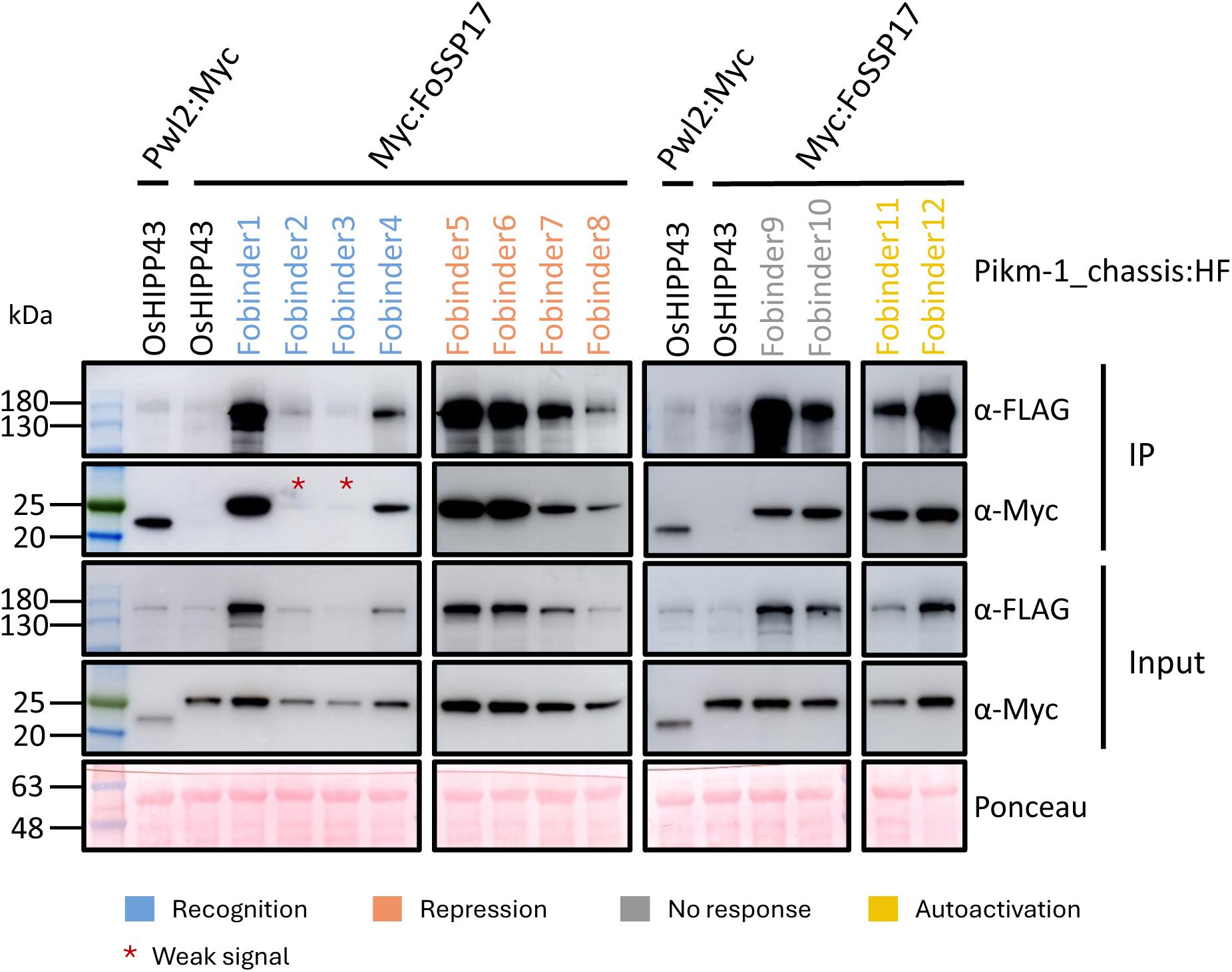
Pikm-1_Fobinder chimeras associate with FoSSP17 in planta. FLAG-tagged Pikm-1_Fobinder chimeras, grouped by their in planta phenotype (Recognition: Fobinder1-4, blue; Repression: Fobinder5-8, salmon; No response: Fobinder9-10, grey; Autoactivation: Fobinder11-12, yellow), were transiently co-expressed with Myc:FoSSP17 in *Nicotiana benthamiana* plants. Co-expression of Pikm-1_HIPP43:HF with Myc:FoSSP17 or Pwl2:Myc were used as negative and positive controls, respectively, alongside each group. At 40 hours post infiltration, leaf samples were harvested and proteins extracted. Proteins in both input and immunoprecipitated (IP) samples were detected by immunoblotting using anti-FLAG and anti-Myc antibodies. Anti-FLAG beads were used for immunoprecipitation; association between all twelve Pikm-1_Fobinders and FoSSP17 was detected, although signal intensities varied among combinations. Asterisks denote the weaker FoSSP17 signals observed for Fobinder2 and Fobinder3, likely due to cell death occurring prior to sample harvest. Similar protein loading was confirmed by Ponceau S staining. The experiment was performed three times with similar results.

### Analytical SEC indicates Fobinder designs form stable complexes with FoSSP17 in vitro

The observation that Fobinders can associate with FoSSP17 in planta to trigger cell death suggests these designed integrated domains are interacting with the effector, consistent with their initial design. To confirm this, we embarked on structural and biophysical characterisation of the Fobinder/FoSSP17 complexes in vitro. Both effector and designs were expressed via recombinant expression in *Escherichia coli* and purified via immobilised metal affinity chromatography (IMAC) coupled with size-exclusion chromatography (SEC) to purify to homogeneity for downstream analyses (Table S1). Of the twelve Fobinders, Fobinder6 failed to purify as it was not stable after IMAC purification.

To determine whether Fobinder designs associate with FoSSP17 to form complexes in solution, equimolar mixtures of each Fobinder with FoSSP17 were subject to analytical SEC on a Superdex S75 5/150 increase GL column, with elution fractions analysed by SDS-PAGE (Figure 5 A; Figure S5). The interaction of Fobinder1 with FoSSP17 is presented as a representative example (Figure 5 A). When mixed, FoSSP17 and Fobinder1 produced a peak at an earlier elution volume relative to FoSSP17 alone (Fobinder1 lacks absorbance at 280 nm), supporting complex formation. SDS-PAGE analysis of co-eluted fractions confirmed the presence of both FoSSP17 (∼17 kDa) and Fobinder1 (∼12 kDa) in the shifted peak (Figure 5 A). Of the eleven Fobinder designs that could be expressed as stable, soluble proteins in *E. coli*, Fobinders 1, 5, and 9 produced a co-elution peak consistent with stable complex formation with FoSSP17 (Figure S5). The remaining eight designs did not co-elute with FoSSP17 under these conditions, including Fobinder4 despite associating with FoSSP17 in planta via co-IP and demonstrating the ability to trigger effector-dependent cell death (Figure 5 A).

**Figure 5.**
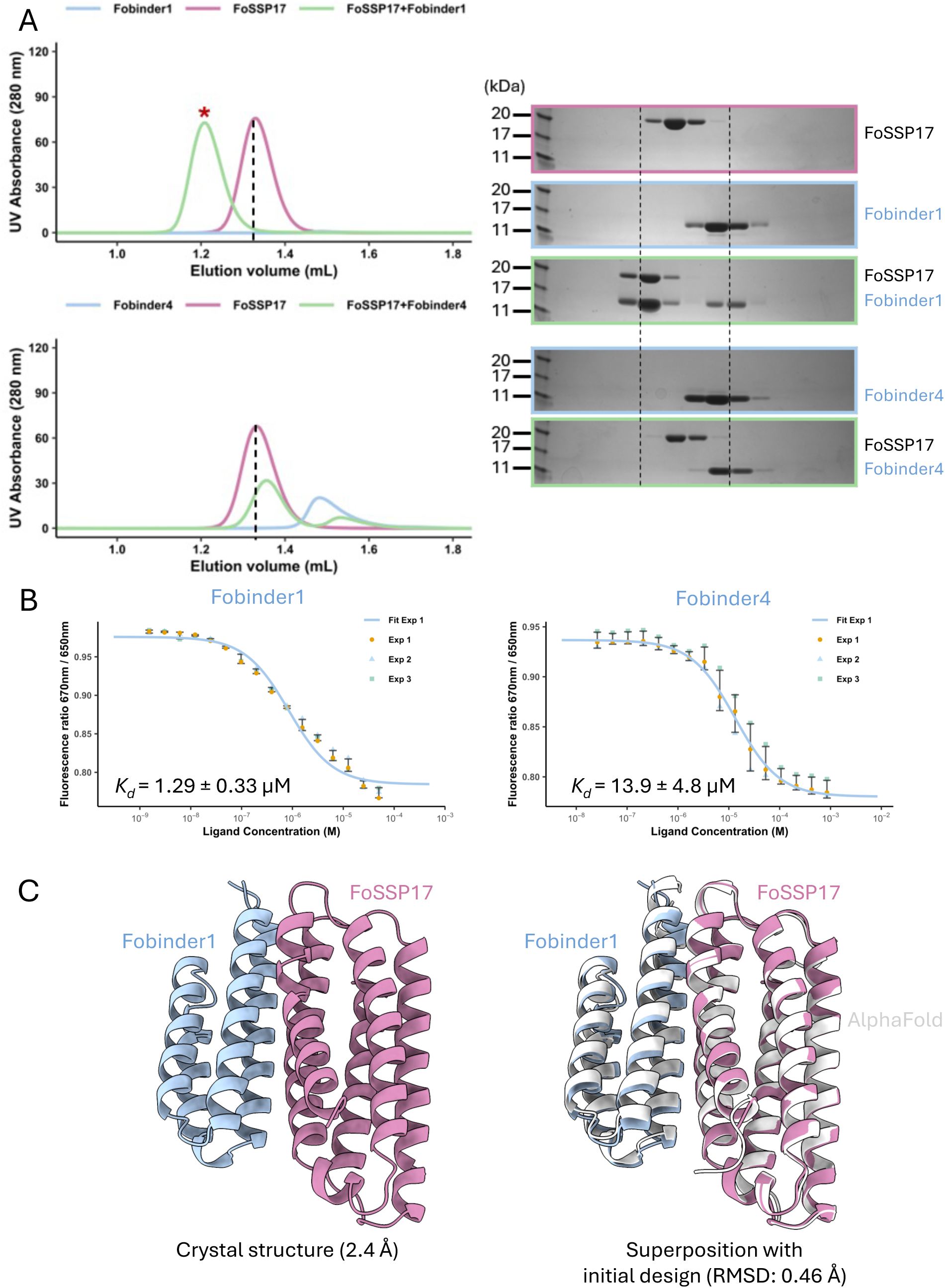
Crystal structures and binding kinetics of FoSSP17-targeting Fobinders demonstrate high fidelity to the design prediction. (A) Fobinder1, but not Fobinder4, forms a stable complex with FoSSP17 in analytical size-exclusion chromatography (SEC) assays. Analytical SEC chromatograms of FoSSP17, Fobinder1, and the FoSSP17/Fobinder1 mixture (top), and of FoSSP17, Fobinder4, and the FoSSP17/Fobinder4 mixture (bottom), are shown on the left. SDS-PAGE gels on the right show the corresponding elution fractions. The elution volume of FoSSP17 alone is indicated by a dashed line; the red asterisk marks the earlier-eluting peak corresponding to FoSSP17/Fobinder1 complex formation. (B) Spectral shift assays measuring the binding of FoSSP17 to Fobinder1 (K*_d_* = 1.29 ± 0.33 μM) and Fobinder4 (K*_d_* = 13.9 ± 4.8 μM). Data from three independent experiments are shown; the fitted curve from one representative experiment is plotted for each. (C) The crystal structure of the Fobinder1/FoSSP17 complex, resolved at 2.4 Å, and its superposition with the AlphaFold2-predicted model of the original RFdiffusion design (RMSD = 0.46 Å), demonstrating high structural similarity between the predicted and experimentally determined complex.

### Spectral shift assays reveal a range of binding affinities across the Fobinder panel

To quantify Fobinder/FoSSP17 binding affinities, spectral shift assays were performed on a panel of five designs (Fobinder1, 4, 5, 9 and 12) spanning the observed range of in planta phenotypes: recognition, no response, repression and autoactivity, respectively (Figure 5 B; Figure S6). Binding was detectable for all five designs, yielding mean dissociation constants (K*_d_*) values ranging from 0.484 ± 0.03 μM to 60.9 ± 14.60 μM across designs from three independent replicates (Figure S6). For Fobinders capable of recognising FoSSP17 in planta, we measured a K*_d_* value of 1.29 ± 0.33 μM for Fobinder1 and 13.9 ± 4.80 μM for Fobinder4 (Figure 5 B). Interestingly, we found binding affinities between Fobinders did not correlate with recognition phenotypes in planta, with the strongest affinity observed being between Fobinder5 and FoSSP17 (0.484 ± 0.03 μM), which was associated with FoSSP17-dependent repression of autoactivity (Figure S6). We also noted Fobinder4 had moderate affinity for FoSSP17 (13.9 ± 4.80 μM) despite not forming a complex in analytical SEC, suggesting transient association between the two proteins in vitro (Figure 5 B; Figure S6).

### The crystal structure of FoSSP17 reveals a homodimeric interface that is maintained in solution

To validate Fobinders were interacting with FoSSP17 as designed with RFdiffusion, purified complexes of Fobinders and FoSSP17 were subject to sparse matrix crystal screening for structure determination via X-ray diffraction. Crystals were obtained for FoSSP17 complexes with Fobinder1, 5 and 9 (Figure S7), subjected to X-ray diffraction at the Diamond Light Source (Oxford, UK), with data collected at a resolution of 2.4 Å, 4.0 Å and 2.0 Å, respectively. Additionally, we crystallised the FoSSP17 effector alone, and collected a dataset at a resolution of 1.75 Å, as no prior structural information existed for this effector.

Structural elucidation of the FoSSP17 effector revealed an amphipathic five-helix bundle fold (helices designated ⍺1-5) consistent with AlphaFold2 predictions, with the experimental structure and prediction having an RMSD of 0.45 Å upon superimposition (Figure S8). Unexpectedly, we observed a homodimer in the FoSSP17 crystal structure with density corresponding to triethylene glycol bound at the core of each monomer. The FoSSP17 dimer has a symmetrical interface formed primarily through hydrophobic interactions of residues of the ⍺2 and ⍺4 helices, supported by a hydrogen bonding network between Arg-72 and His-65 with Asp-132 at the C-terminal ends of ⍺2 and ⍺4 (Figure S8).

To test whether FoSSP17 forms a dimer in solution, we subjected purified FoSSP17 to size-exclusion chromatography coupled with right-angle laser light scattering (SEC-RALS) using an OMNISEC system (Malvern Panalytical) to determine the molecular weight in solution. We measured an average molecular weight of 34.1 kDa across the FoSSP17 elution peak, consistent with approximately double the predicted monomeric mass of the effector (17.2 kDa), indicating FoSSP17 forms a dimer in solution (Figure S9).

### Structure determination of Fobinder/FoSSP17 complexes validates computational binder design

In total, we were able to obtain three crystal structures of Fobinders in complex with FoSSP17 that resulted in a variety of different in planta phenotypes, these being Fobinder1 that mediated recognition of the effector (Figure 5; Figure S10), Fobinder5 which demonstrated effector-dependent cell death repression (Figure S10), and Fobinder9 which did not result in a cell death response despite mediating association of FoSSP17 with the Pik-1 receptor in planta (Figure S10, Figure S11). Upon superposition of the binder/effector complexes, it was apparent that Fobinder1, 5 and 9 were each able to form a complex with FoSSP17 through a similar interface consisting of residues of the ⍺3 and ⍺5 helices of the effector (Figure S10). We would note that the Fobinder5/FoSSP17 structure was determined from limited-resolution diffraction data, and as such only overall domain positions were placed in the electron density maps without further refinement or model rebuilding.

Analysis of the Fobinder1/FoSSP17 complex through superposition of the initial RFdiffusion design and experimentally determined complex demonstrated high structural homology (RMSD = 0.46 Å; Figure 5 C, Figure S11), with the effector binding at our designed protein interaction interface with side-chain level accuracy (Figure S12). These data confirm that binding and recognition of FoSSP17 by the Pik-1 receptor incorporating Fobinder1 results from the molecular contacts specifically designed with RFdiffusion.

Design of Fobinders with RFdiffusion did not account for homodimer formation of FoSSP17, using a monomer of FoSSP17 predicted with AlphaFold (Figure S1) as the design target input because no prior structural or biochemical information existed for this effector prior to this study. As such, we sought to understand the potential impact of the FoSSP17 homodimer on the interaction with different Fobinders. Superposition of the AlphaFold2-predicted Fobinder/FoSSP17 designs onto the FoSSP17 homodimer suggested that dimerisation may partially occlude the predicted binding interface for several Fobinders (Figure S13). This observation may account for the absence of a co-elution peak for these designs in analytical SEC, but despite this, we still observed association with the Pikm-1_Fobinder chimeras in co-immunoprecipitation assays (Figure 4). By contrast, Fobinders 1, 5, and 9 are predicted to engage a surface not overlapping with the dimer interface, consistent with the in vitro binding observed for these designs (Figure 5; Figure S6). Notably, Fobinder4 produced effector-dependent cell death in planta despite the absence of a co-elution peak in analytical SEC, presenting conflicting observations not fully explained by dimer-interface occlusion alone. Spectral shift assays measured a K*_d_* of approximately 14 μM for Fobinder4 (Figure 5B), indicating that a weak interaction with FoSSP17 is detectable in vitro, and co-immunoprecipitation assays confirmed in planta association between Fobinder4 and FoSSP17 (Figure 4). These data raise the possibility that Fobinder4 engages FoSSP17 transiently in vitro at a level insufficient to shift the elution profile in analytical SEC, yet sufficient to modulate NLR signalling in the cellular context.

## Discussion

Historically, developing bespoke disease resistance in plants through NLR engineering has been a labour-intensive process, reliant on discovery and extensive characterisation of natural resistance genes or detailed molecular dissection of plant-pathogen interactions^36–38^. Bioengineering approaches that integrate foreign effector binding domains into NLR receptors, including host derived HMA domains^3,26,27,39,40^ and VHH nanobody grafting^3,25^, have previously demonstrated the ability to directly manipulate effector recognition profiles of NLRs, but in each case these approaches required prior effector characterisation or, in the case of nanobodies, an immunisation campaign, which significantly limits throughput. The Pik system has previously been engineered to recognise effectors from *M. oryzae* outside its native recognition profile^3,23,26,27^, but our work extends this capacity to FoSSP17, an effector from *F. oxysporum* f. sp*. cubense*^28^, a pathogen of banana with no evolutionary or host relationship to the rice-*M. oryzae* pathosystem from which Pik-1 is derived. Critically, this proof-of-concept manipulation of Pik-1 to recognise FoSSP17 in *N. benthamiana* was performed without the need for extensive prior characterisation of the effector structure or function.

### De novo protein binders against effectors can be generated with high fidelity

At the onset of this study, we performed protein design campaigns with RFdiffusion^7^ against three effectors; two from *M. oryzae*, AVR-PikF and AVR-Pii, and one from *Foc* TR4, FoSSP17. As we filtered these designs through *N. benthamiana* for activity, it is not possible to say, and assumedly unlikely, that all binders we generated maintained the ability to bind to their cognate effector targets as per their initial design. However, for the Fobinders that we did take through structural and biophysical characterisation (Fobinders1, 5, 9; Figure 5 C; Figure S10; Figure S11), experimentally determined structures in complex with FoSSP17 show that each protein engages the effector through the interface indicated by the RFdiffusion design (Figure 5 C; Figure S10; Figure S11). Indeed, if we take Fobinder1 as an example, we demonstrate that we can design a specific interaction with an effector with side chain level accuracy that results in recognition upon integration into Pikm-1 (Figure 2; Figure 5; Figure S12), further evidenced by superposition of the experimentally determined Fobinder1/FoSSP17 complex with the AlphaFold prediction of the initial design with an RMSD of 0.46 Å.

Together, these results indicate that RFdiffusion^7^ and ProteinMPNN^6^ can generate binders against plant pathogen effectors with high structural fidelity, including against effectors for which no experimental structure exists, as was the case for FoSSP17 (Figure S1). However, the three Fobinder/FoSSP17 structures show intrinsically similar binding modes, yet span three distinct in planta outcomes upon integration into Pikm-1: recognition, repression, and no response (Figure S10). Therefore, while we can be confident that the designed binding event is reproducible and accurately predicted, other factors outside of effector binding clearly influence variability in the functional outcome.

### Signal relay is the most significant bottleneck to engineering expanded recognition in NLRs

The central finding of this study is not simply that recognition can be engineered, but that recognition depends on two separable engineering problems, effector binding and signal relay. Co-immunoprecipitation assays demonstrate that all twelve Pikm-1_Fobinder chimeras tested associate with FoSSP17 upon co-expression in planta (Figure 4), regardless of their downstream cell death phenotype, with only a small subset of these interactions translating into effector-dependent cell death (Figure 2, Figure S2). As such, we argue that the principal bottleneck for NLR engineering with de novo binders is not the design of binding itself, which is well supported by our structural and biophysical data (Figure 5), but signal relay of that binding event into immune receptor activation. Indeed, of the 44 AVR-PikF binders tested, 30 produced effector-independent cell death upon integration into Pikm-1, and a comparable proportion was observed for the FoSSP17 panel, in which 31 of 46 Fobinder chimeras were autoactive in the absence of effector (Figure S2).

This pattern echoes previous bioengineering efforts in the Pik system, where integration of nanobody, host-derived HMA, or other foreign domains into Pik-1 has repeatedly caused autoactivation^3,25–27^. Furthermore, the disconnect between effector binding and signal relay is well illustrated here by absence of any consistent relationship between binding strength and functional outcome. Fobinder5, for instance, bound FoSSP17 with the highest affinity measured across the panel (K*_d_* = 0.48 μM) yet produced a repression phenotype rather than activation, while Fobinder4 bound with substantially lower affinity (K*_d_* = 13.90 μM) showing no detectable complex by analytical size exclusion chromatography, yet reliably triggered effector-dependent cell death and precipitated the effector in planta (Figure 5 B, Figure S6), though the lower affinity of Fobinder4 for FoSSP17 may also be an artifact of competing for the same binding interface as the FoSSP17 homodimer (Figure S8; Figure S13). If Pik activation was governed primarily by the strength of the binding event, we would expect a correlation between binding affinity and cell death strength in *N. benthamiana*. As we do not observe this, effector recognition through binder integration therefore likely depends on additional properties of the binder beyond interaction with the effector.

Unfortunately, due to a lack of structural and mechanistic understanding of how the integrated domain of the Pik-1 receptor translates effector binding into an activation signal for the rest of the receptor, there exists no comparable design input for engineering signal relay within the NLR. This underscores the need for pre– and post-activation structures of a Pik resistosome to aid in guiding the design of signal relay in these receptors in future bioengineering efforts.

### FoSSP17-dependent repression of autoactive Pik-1_Fobinder chimeras suggests avenues for NLR disruption by plant pathogen effectors

The observation of effector-dependent repression was an unexpected phenotype, but offers insight into how NLR signalling may be disrupted by a pathogen-derived molecule. Fobinders 5-8, when integrated into Pikm-1 and co-expressed with Pikp-2, produced autoactive cell death in the absence of the effector. Co-expression with FoSSP17 attenuated this autoactivity (Figure 3 A), and this attenuation was effector-specific: co-expression with the unrelated effectors AVR-PikD or Pwl2 did not reproduce the repression phenotype (Figure 3 A and C), and the autoactive Pik pair Pi1-5C/Pi1-6C, remained constitutively active in the presence of FoSSP17 (Figure 3 B and C), suggesting repression of cell death is not an innate function of FoSSP17. Repression depends therefore on a specific interaction between the effector and Fobinder domain itself, rather than a general suppressive effect of FoSSP17 on Pik-mediated signalling.

This observation of FoSSP17-dependent cell death repression is consistent with one of the two following scenarios: 1) that effector binding holds the chimera in an inactive conformation as it assembles, without permitting the transition to a signalling-competent state, or 2) an already-active, signalling-competent NLR complex may be disassembled or otherwise disarmed by a bound effector post-activation. If the latter is true, this would imply a more dynamic model of NLR regulation than generally assumed, one where activation may not be a one-way switch but a state that pathogens might have evolved to intervene in order to reverse. Irrespective of either scenario, the repression phenotype observed here is consistent with the broader precedent of effector-mediated suppression of NLR activity^41–44^, as best described by the recent cryo-EM structures of the *Phytophthora infestans* effector AVRcap1B bound to a nascent tomato NRC3 resistosome^41^, preventing its full assembly into a signalling-competent resistosome.

Due to the simultaneous co-expression of effector and NLR used in our cell death assays, we cannot delineate whether the effector is preventing activation or reversing it once it has already occurred, since receptor and effector are present together from the outset. To understand this, a temporally controlled or inducible effector delivery system, paired with a biochemical readout of receptor oligomeric state before and after effector exposure, would be needed.

### Protein design as a tool for understanding NLR biology

Beyond its translational aim, this study points to an unanticipated use for de novo protein design: as a tool for isolating variables that are otherwise difficult to separate in natural NLR variation. As previously discussed, Fobinder1, Fobinder5 and Fobinder9 bind FoSSP17 through essentially the same designed interface, as confirmed by crystal structures with consistent geometry across the panel (Figure 5 C; Figure S10; Figure S11), and yet produce three distinct outcomes when integrated into Pikm-1 and challenged with the effector: recognition, repression and no response, respectively. This is a separation of variables that is difficult to achieve using naturally occurring allelic diversity, where polymorphisms typically accumulate across the integrated domain as a result of co-evolution with the effector and are rarely confined to a single, structurally defined interface^44–47^. This may offer a route to dissecting determinants of NLR activation that is largely orthogonal to existing approaches, which are based on natural variants or targeted point mutagenesis. Where prior studies of Pik incompatibility have relied on chimeras between natural alleles or systematic mutagenesis of the integrated domain to identify regions implicated in signal relay^3,24,46^, a structurally confirmed, phenotypically diverse binder panel offers a complementary resource in which the contribution of binding geometry has already been controlled. Used in this way, de novo design functions not only as an engineering strategy for generating new recognition specificities, but as an experimental system for asking which features of an integrated domain, beyond its capacity to bind, govern signal relay into NLR receptor activation.

The repression phenotype discussed previously is itself an example of this orthogonal value in probing NLR biology through design. The question raised by this observation, whether a pathogen-derived molecule can disrupt an NLR complex after activation rather than only before it, is not one that natural allelic variation or conventional mutagenesis is well positioned to probe directly, since both approaches act on the receptor rather than supplying a defined, structurally validated ligand whose timing and binding mode are independently known. A designed binder panel, by contrast, offers a starting point from which binding, signal relay and the resulting recognition outcome can, in principle, be interrogated as separable layers of the same problem, rather than inferred jointly from a single phenotype.

### Conclusions

This work demonstrates a computationally designed, de novo protein domain can function as an integrated sensory module within a plant immune receptor, establishing protein design as a viable strategy for plant immune engineering. This advance does not remove the co-evolutionary pressure that pathogens exert on any effector-based resistance strategy, but it does offer a path to a substantially compressed timescale in which new recognition specificities may be generated and deployed against emerging effector variants, particularly as new computational pipelines such as BindCraft^49^, BoltzGen^50^and RFdiffusion3^51^ continue to improve on protein binder design methods used in this study. Realising this potential in the field will require demonstrating that designed receptors retain immune function in stably transformed crops, a translation that for *Foc* TR4 carries the additional burden of effective resistance in root tissue against a soil-borne pathogen, and of stable transformation in banana itself. We anticipate that as design pipelines mature and structural understanding of NLR activation deepens in parallel, de novo protein design will become an increasingly central tool, both for engineering bespoke disease resistance in crops and for probing the fundamental mechanisms behind immune receptor activation. The plug-and-play modularity demonstrated here for Pik-1, together with the prospect of extending these principles to other NLRs and to extracellular receptors such as PRRs and RLKs, points to a future in which plant immunity can be engineered with a precision and speed not previously available to the field.

## Supplemental Figure Legends

**Figure S1.**
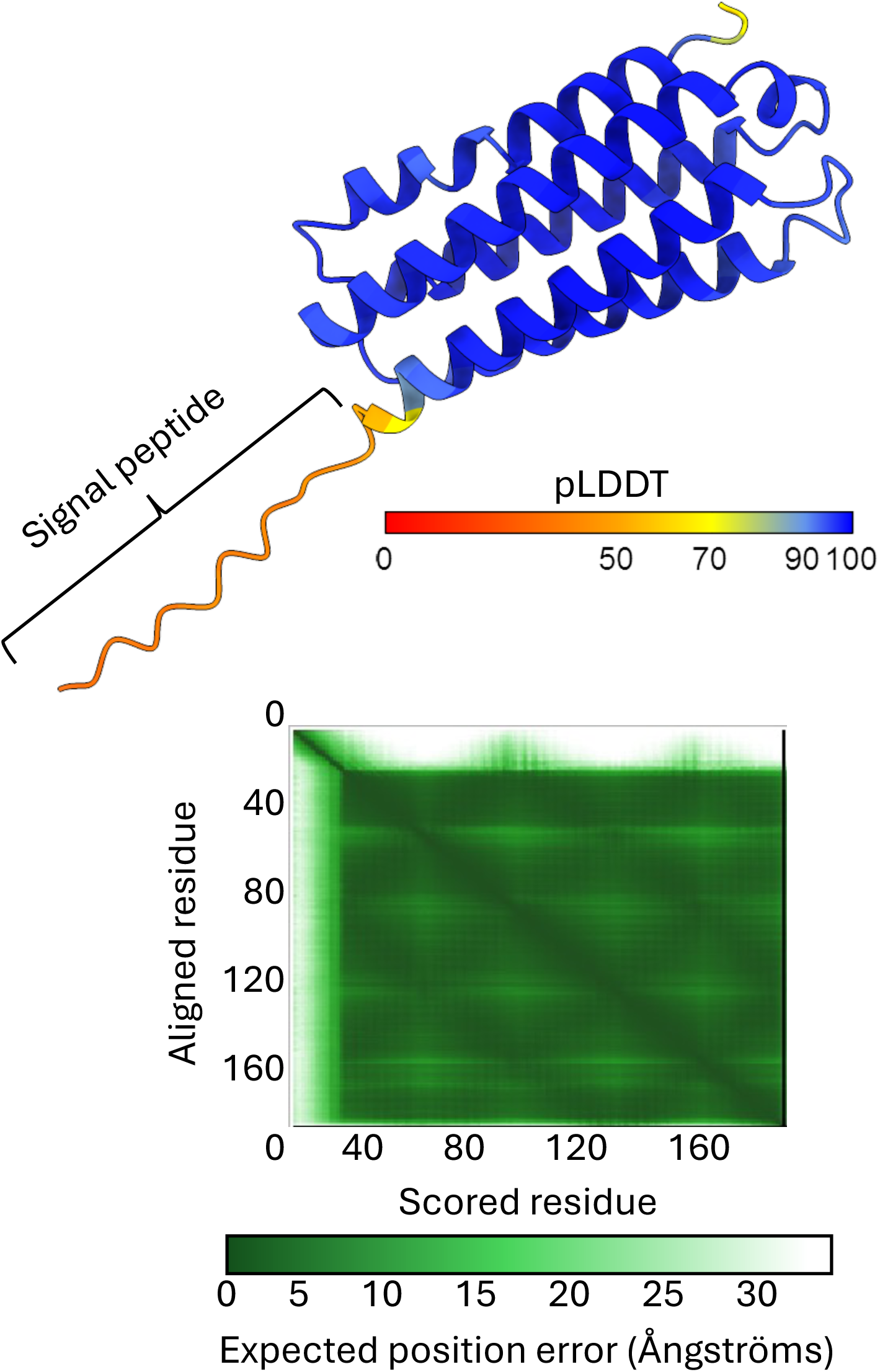
Prediction of the FoSSP17 effector with AlphaFold2. The structure of FoSSP17 was predicted with AlphaFold2 v1.5.5 as implemented in ColabFold^33,34^. **Top:** predicted model coloured by per-residue confidence (pLDDT); the prediction was high confidence overall (average pLDDT = 90.8, pTM = 0.865), with disorder confined to the predicted signal peptide of the effector. **Bottom:** predicted aligned error (PAE) plot, indicating high confidence in the relative positioning of residues across the modelled structure.

**Figure S2.**
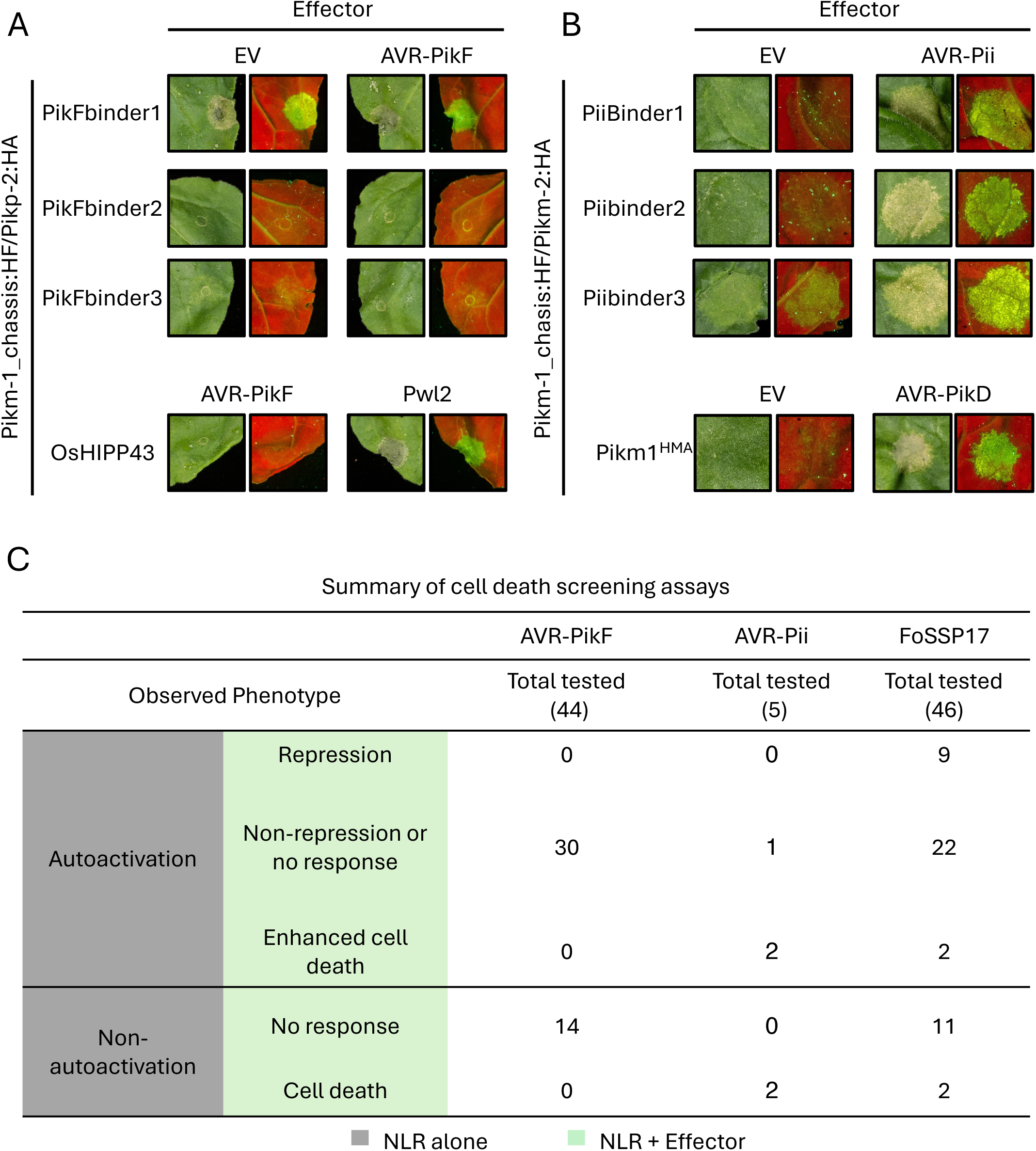
Summary of *Nicotiana benthamiana* cell death screening of Pik-1 binder chimeras against the AVR-PikF, AVR-Pii and FoSSP17 effectors. (A) Representative leaf images from the cell death screening of AVR-PikF binders (PikFbinder1–3). The de novo designed AVR-PikF binders were introduced into the Pikm-1_chassis:HF/Pikp-2:HA background. Co-expressing Pikm-1_OsHIPP43/Pikp-2 with AVR-PikF or Pwl2 served as negative and positive controls, respectively. (B) Representative leaf images from the cell death screening of AVR-Pii binders (PiiBinder1–3), introduced into the Pikm-1_chassis:HF/Pikm-2:HA background. Wild-type Pikm1 co-expressed with empty vector (EV) or AVR-PikD served as negative and positive controls, respectively. (C) Summary table of cell death screening outcomes for AVR-PikF, AVR-Pii and FoSSP17 binders, classified first by whether the Pik-1 binder chimera was autoactive in the absence of effector (NLR alone, grey), and then by the effect of effector co-expression on that phenotype (NLR + Effector, green): repression of autoactivity, non-repression or no change, or enhanced cell death (for autoactive chimeras); and no response or effector-dependent cell death (for non-autoactive chimeras). Total numbers of binders tested per effector are indicated.

**Figure S3.**
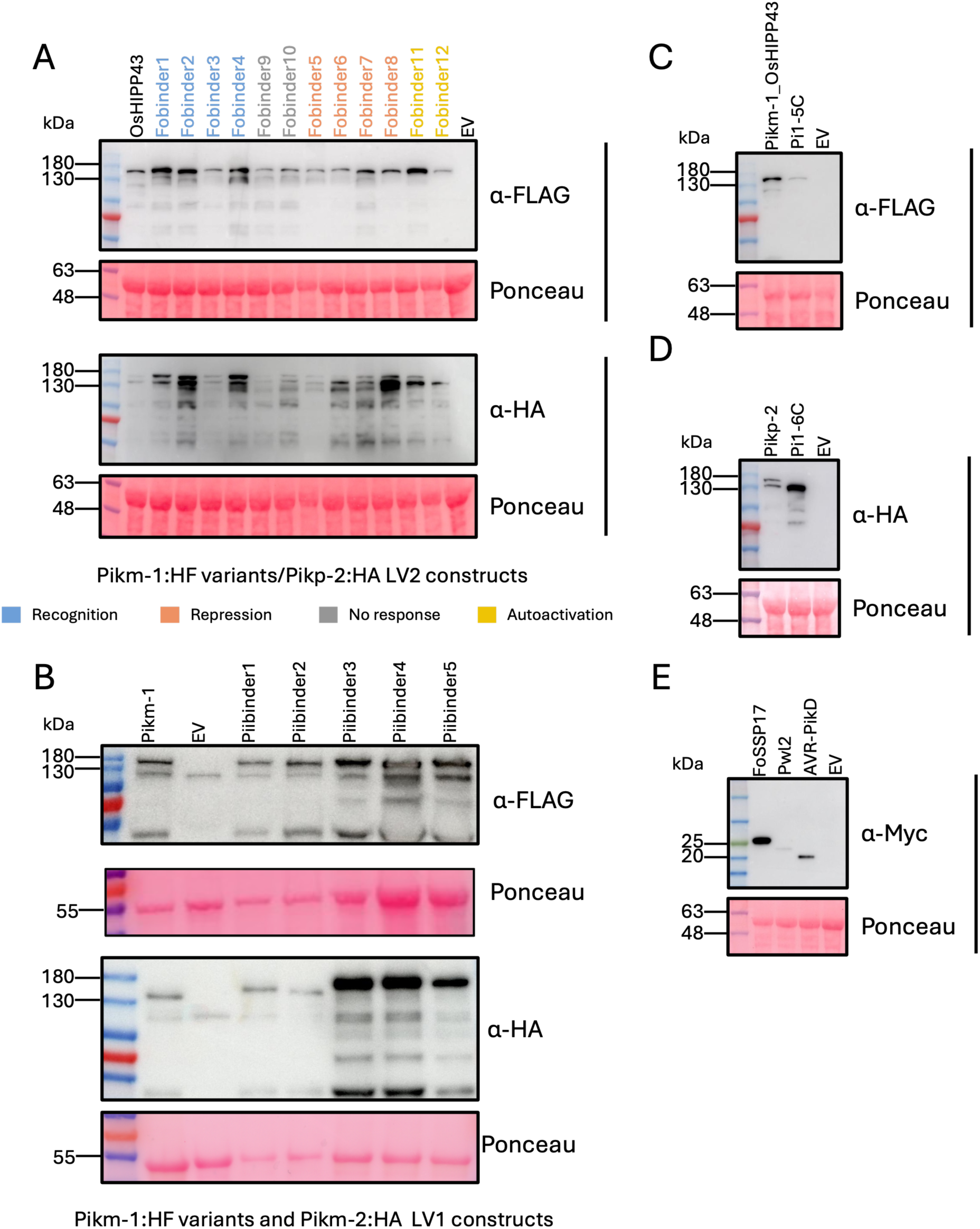
The protein accumulation of Pikm-1_Fobinder chimeras, Pik variants and effectors used in this study in *Nicotiana benthamiana* plants as detected by western blot. (A) Accumulation of FLAG-tagged Pikm-1_OsHIPP43 and Pikm-1_Fobinder chimeras (coloured by phenotype: Fobinder1–4, Recognition, blue; Fobinder9–10, No response, grey; Fobinder5–8, Repression, salmon; Fobinder11–12, Autoactivation, yellow), alongside HA-tagged Pikp-2, in the Pikm-1:HF variants/Pikp-2:HA LV2 background, detected by anti-FLAG and anti-HA immunoblotting, respectively. (B) Accumulation of FLAG-tagged wild-type Pikm-1 and Piibinder1–5 chimeras, alongside HA-tagged Pikm-2, detected by anti-FLAG and anti-HA immunoblotting, respectively. (C) Accumulation of FLAG-tagged Pikm-1_OsHIPP43 and Pi1-5C, detected by anti-FLAG immunoblotting. (D) Accumulation of HA-tagged Pikp-2 and Pi1-6C, detected by anti-HA immunoblotting. (E) Accumulation of Myc-tagged FoSSP17, Pwl2 and AVR-PikD, detected by anti-Myc immunoblotting. Empty vector (EV) was included as a negative control throughout. All proteins were extracted from infiltrated leaf tissue at 40 hours post-infiltration. Protein loading was confirmed by Ponceau S staining throughout. Experiments shown were performed three times with similar results.

**Figure S4.**
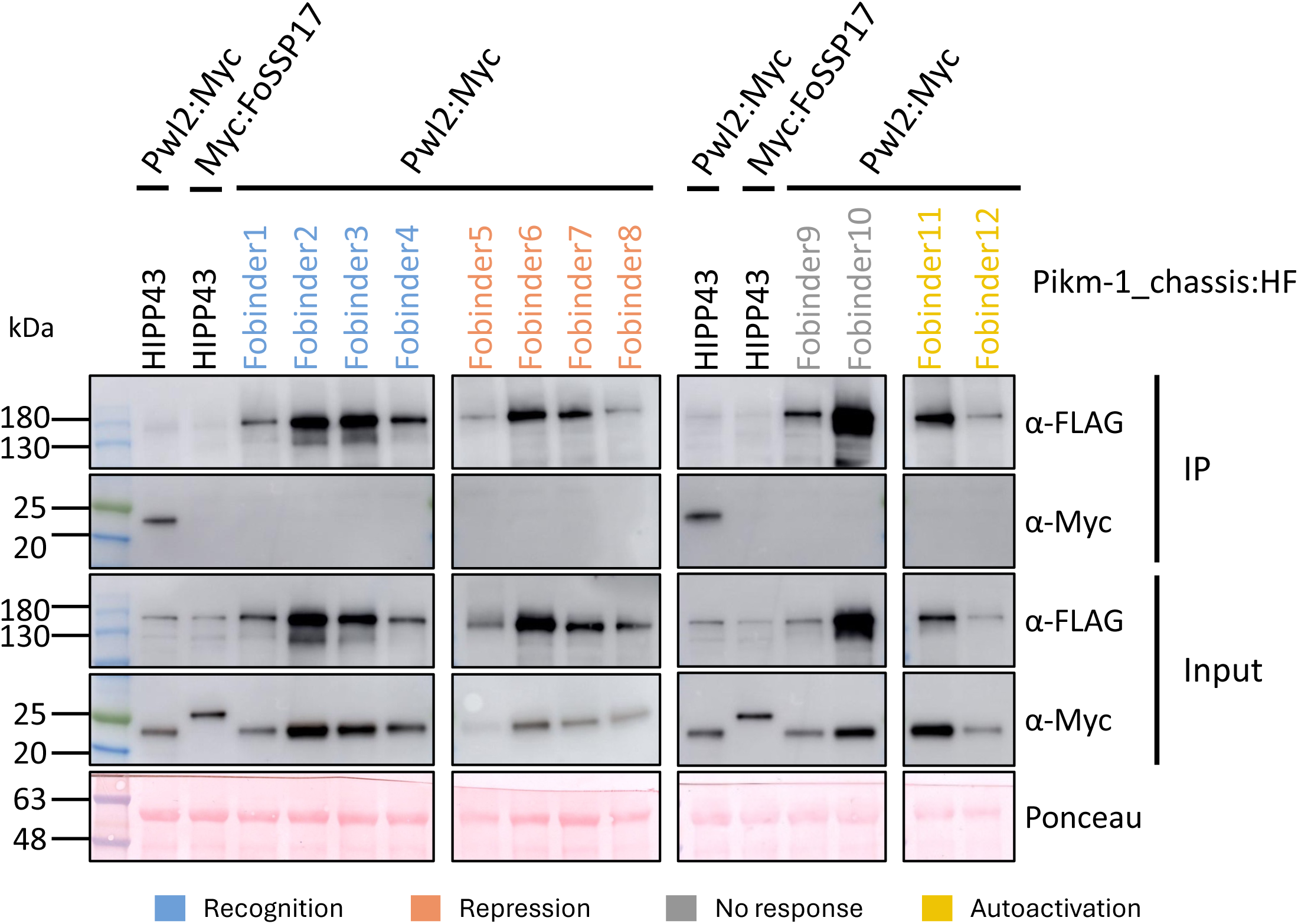
Pikm-1_Fobinder chimeras do not associate with Pwl2 in planta. FLAG-tagged Pikm-1_Fobinder chimeras, grouped by their in planta phenotype (Recognition: Fobinder1-4, blue; Repression: Fobinder5-8, salmon; No response: Fobinder9-10, grey; Autoactivation: Fobinder11-12, yellow), were transiently co-expressed with Pwl2:Myc in *Nicotiana benthamiana* plants. Co-expressing Pikm-1_OsHIPP43:HF with Myc:FoSSP17 and Pwl2:Myc were used as negative and positive controls, respectively, alongside each group. At 40 hours post-infiltration, leaf samples were harvested and proteins extracted. Proteins in both input and immunoprecipitated (IP) samples were detected by immunoblotting using anti-FLAG and anti-Myc antibodies. Anti-FLAG beads were used for immunoprecipitation; none of the twelve Pikm-1_Fobinders showed association with Pwl2, indicating that their interaction with FoSSP17 is specific. Similar protein loading was confirmed by Ponceau S staining. The experiment was performed three times with similar results.

**Figure S5.**
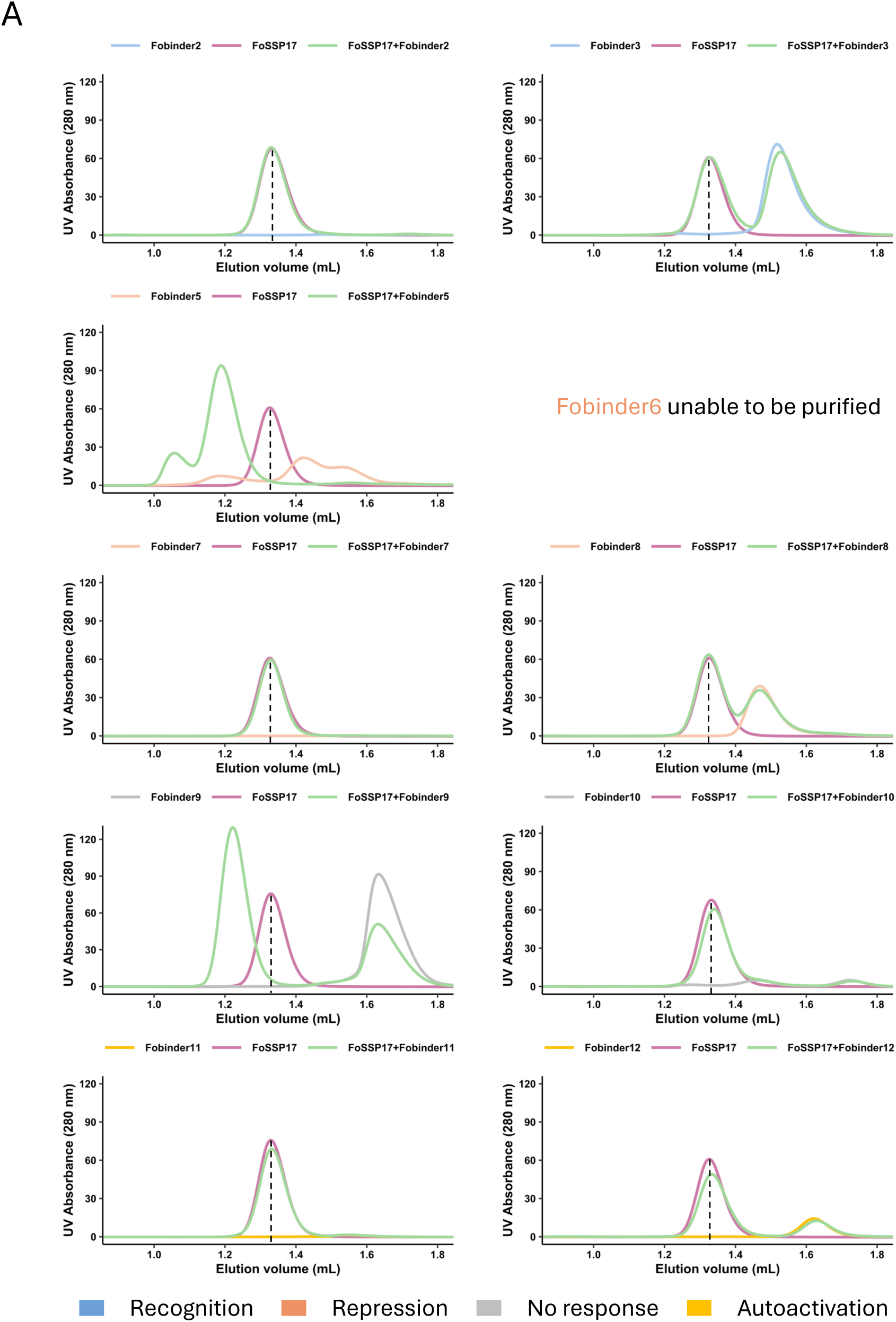
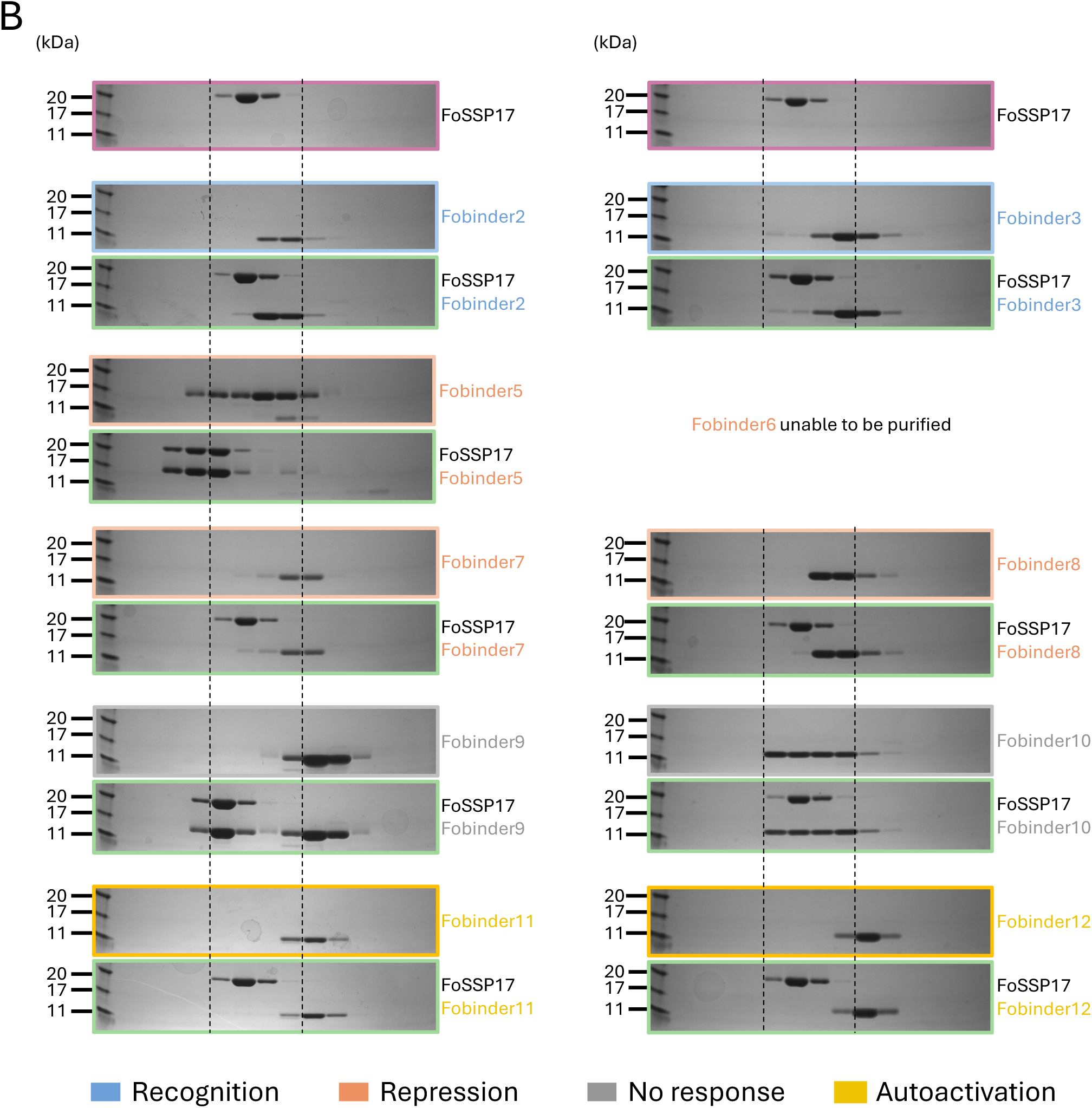
Analytical size exclusion and SDS-PAGE analysis of recombinantly expressed Fobinders with FoSSP17. (A) Analytical SEC traces of FoSSP17 alone, each Fobinder alone, and the FoSSP17/Fobinder mixture, for Fobinder2, 3, 5, 7, 8, 9, 10, 11 and 12 (coloured by phenotype: Recognition, blue; Repression, salmon; No response, grey; Autoactivation, yellow). Fobinder1 and Fobinder4 are shown separately in Figure 5 A. Fobinder6 could not be purified and is excluded from this analysis. The elution volume of FoSSP17 alone is indicated by a dashed line where applicable. (B) SDS-PAGE gels showing the elution fractions corresponding to each trace in (A), confirming the identity of FoSSP17 and each Fobinder by molecular weight.

**Figure S6.**
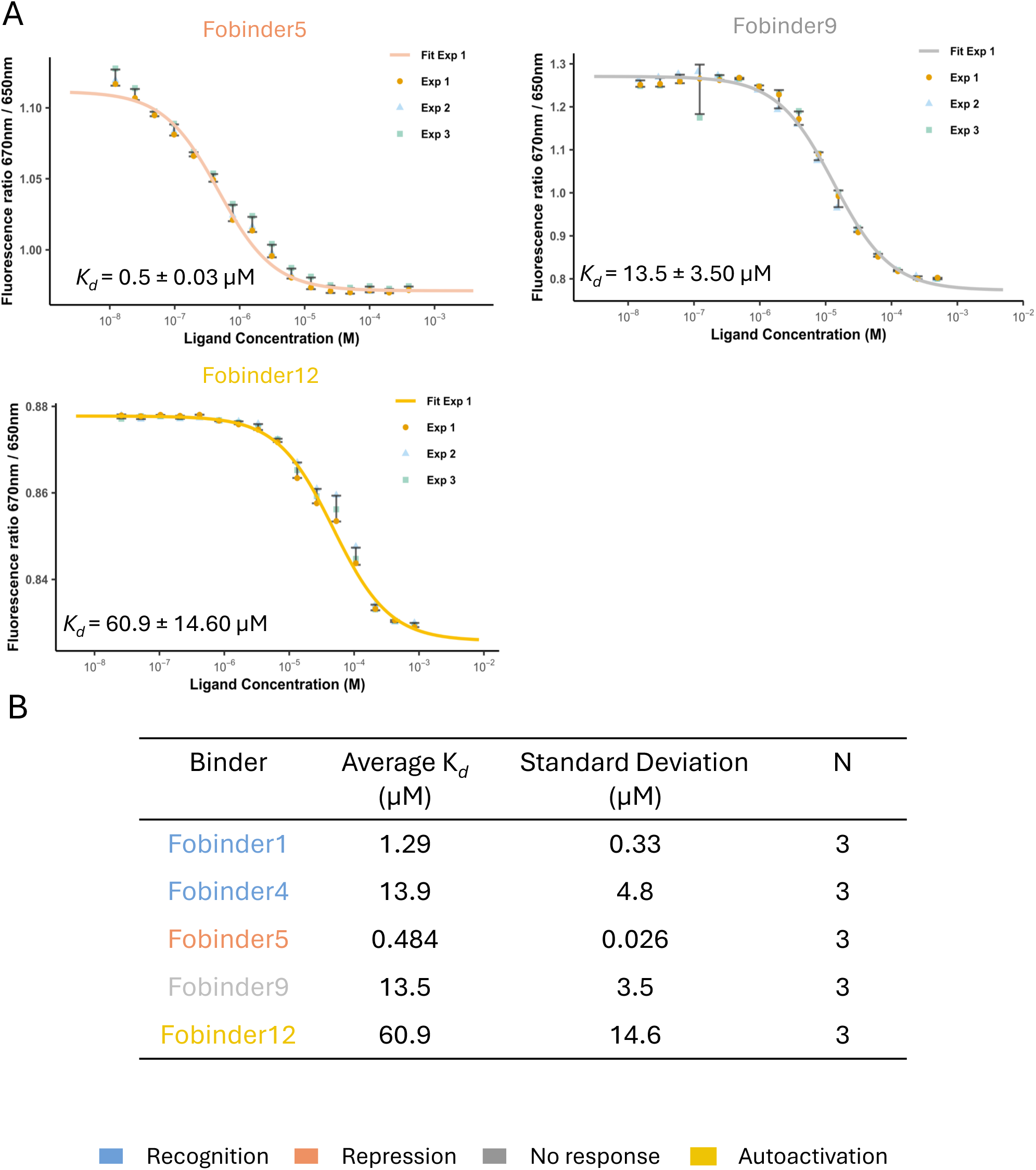
Fobinders across observed in planta phenotypes demonstrate affinity for FoSSP17 in spectral shift assays. (A) Spectral shift assays measuring the binding of FoSSP17 to Fobinder5, Fobinder9 and Fobinder12 (coloured by phenotype: Repression, salmon; No response, grey; Autoactivation, yellow). Data points from three independent experiments are shown for each Fobinder; the fitted binding curve from one representative experiment is plotted, with the average dissociation constant (K*_d_*) indicated. (B) Summary table of binding affinities determined by spectral shift assays for all five Fobinders assayed (Fobinder1, 4, 5, 9 and 12; see also Figure 5 B for Fobinder1 and Fobinder4), coloured by phenotype as in (A). Average dissociation constants (K*_d_*), standard deviations, and the number of independent replicates (N) are shown for each.

**Figure S7.**
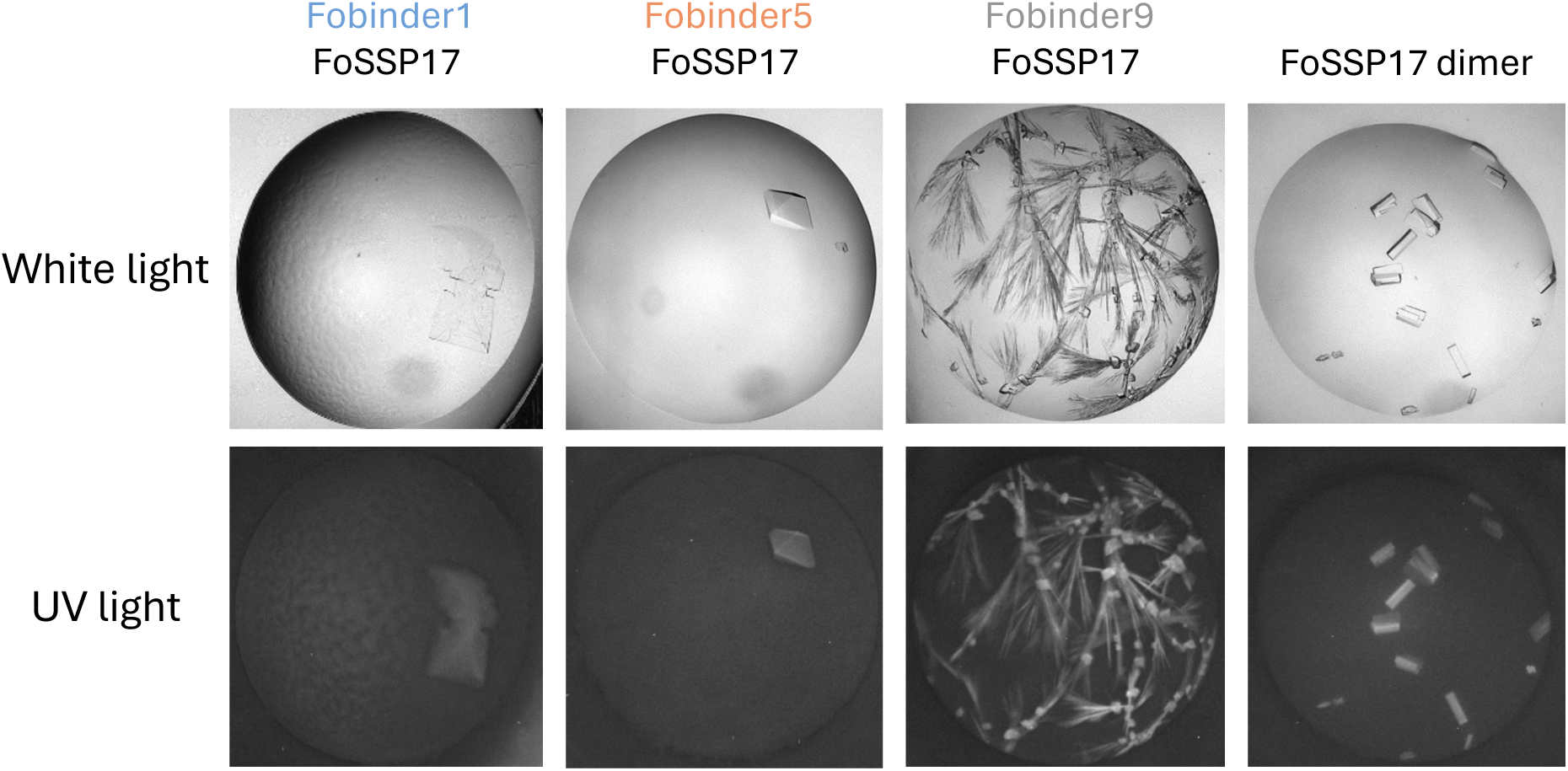
Representative crystals of Fobinder/FoSSP17 complexes and the FoSSP17 dimer obtained by sitting-drop vapour diffusion. Crystals of Fobinder1/FoSSP17 (7 mg/ml; 0.06 M Divalents, 0.1 M Buffer System 1 pH 6.5, 50% v/v Precipitant Mix 2; Morpheus screen, Molecular Dimensions), Fobinder5/FoSSP17 (8 mg/ml; 0.1 M Imidazole pH 7.0, 50% v/v MPD 1; PROPLEX screen, Molecular Dimensions), Fobinder9/FoSSP17 (9.5 mg/ml; 0.06 M Divalents, 0.1 M Buffer System 3 pH 8.5, 50% v/v Precipitant Mix 4; Morpheus screen, Molecular Dimensions) and FoSSP17 dimer (11 mg/ml; 0.1 M Carboxylic acids, 0.1 M Buffer System 3 pH 8.5, 50% v/v Precipitant Mix 1; Morpheus screen, Molecular Dimensions) are shown under white light (top) and UV light (bottom). UV fluorescence confirms the proteinaceous nature of the crystals.

**Figure S8.**
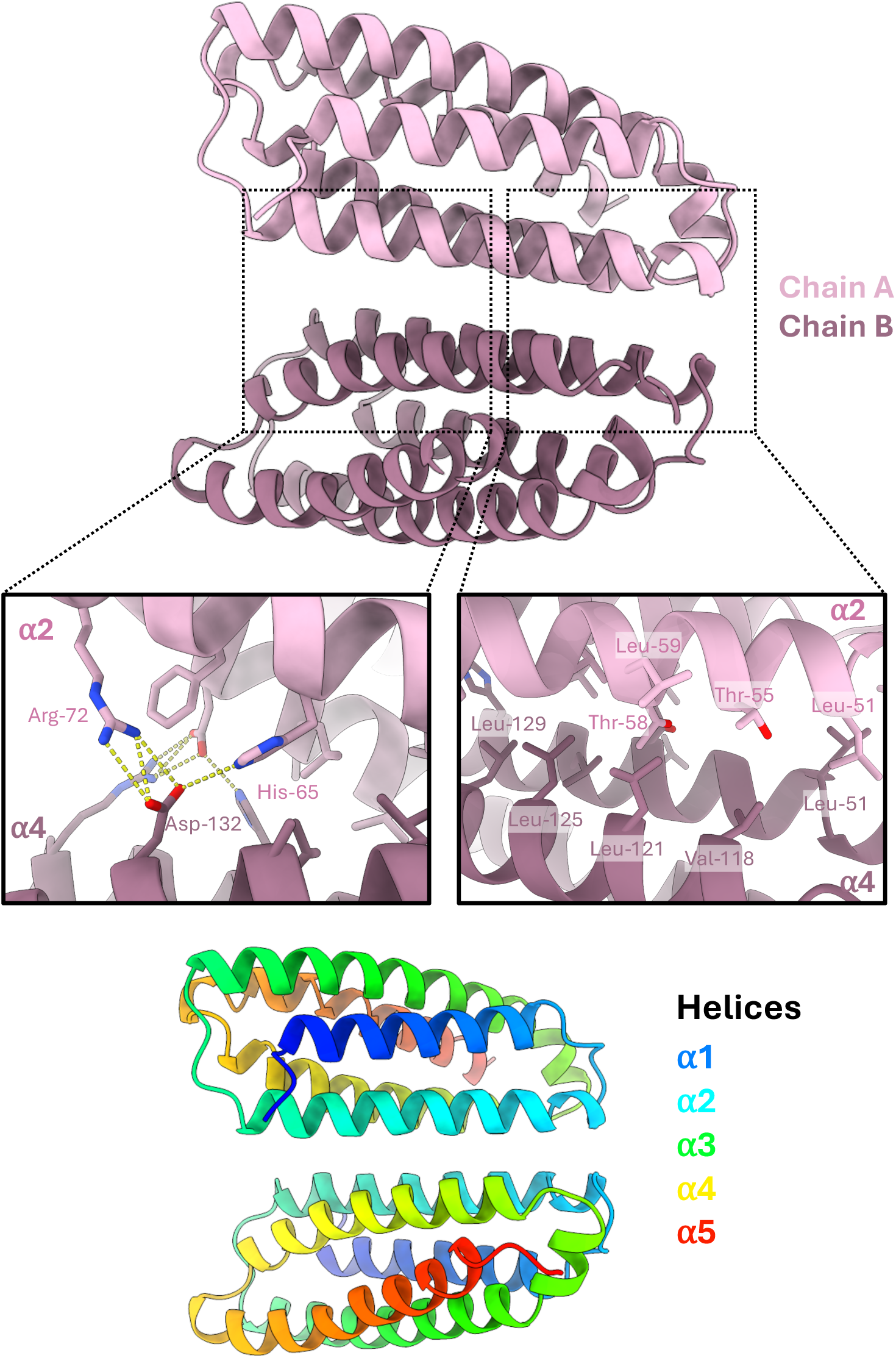
The crystal structure of FoSSP17 reveals a homodimeric interface. Top: the FoSSP17 homodimer crystal structure, with the two monomers labelled Chain A and Chain B. Two inset panels show detail of the dimer interface: (left) the hydrogen bonding and salt bridge network formed by His-65, Arg-72, and Asp-132 of each monomer; (right) the hydrophobic core formed by reciprocal interactions between Leu-51, Thr-55, Leu-59, Val-118, Leu-121, Leu-125, and Leu-129. **Bottom:** a single FoSSP17 monomer coloured by position along the polypeptide chain (rainbow, blue–red, N-terminus to C-terminus), illustrating the arrangement of the five helices (α1–α5) that contribute to the dimer interface shown above.

**Figure S9.**
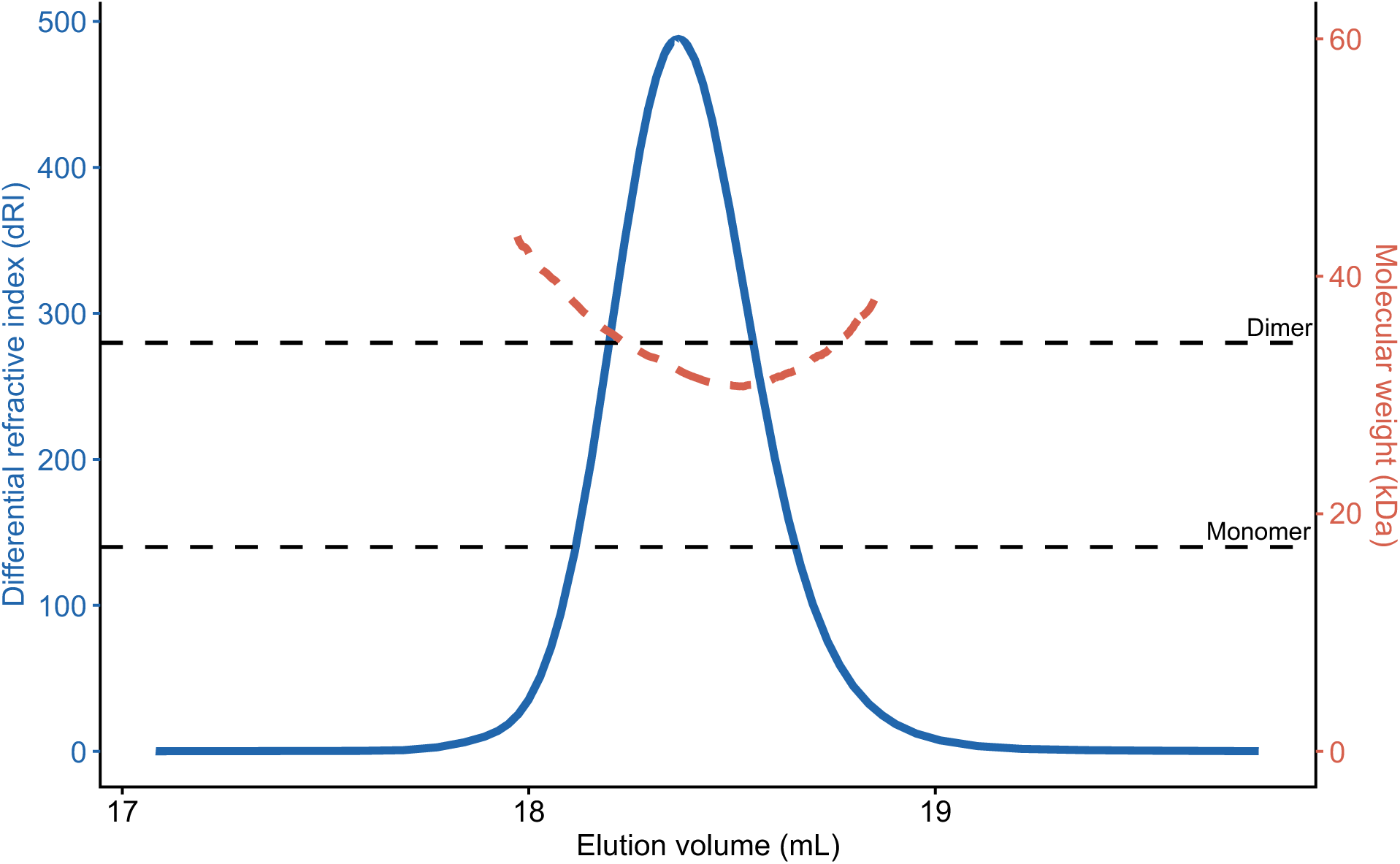
Size exclusion coupled with right angle and low angle light scattering indicates FoSSP17 homodimer in solution. 20 μL purified FoSSP17 at a concentration of 500 μM was injected and separated on a Superose 6 10/300 Increase column and subjected to right-angle and low-angle light scattering (RALS/LALS) with a Malvern Panalytical OMNISEC system. The differential refractive index (dRI, left axis) shows a single elution peak corresponding to FoSSP17. Molecular weight, calculated across the elution peak from the light scattering data (right axis), is shown alongside dashed reference lines indicating the expected molecular weight of FoSSP17 monomer (17.2 kDa) and dimer (34.4 kDa). The calculated molecular weight across the peak closely tracks the dimer reference line, with an average mass of 34.1 kDa, approximately twice the predicted monomeric mass, indicating that FoSSP17 is a dimer in solution.

**Figure S10.**
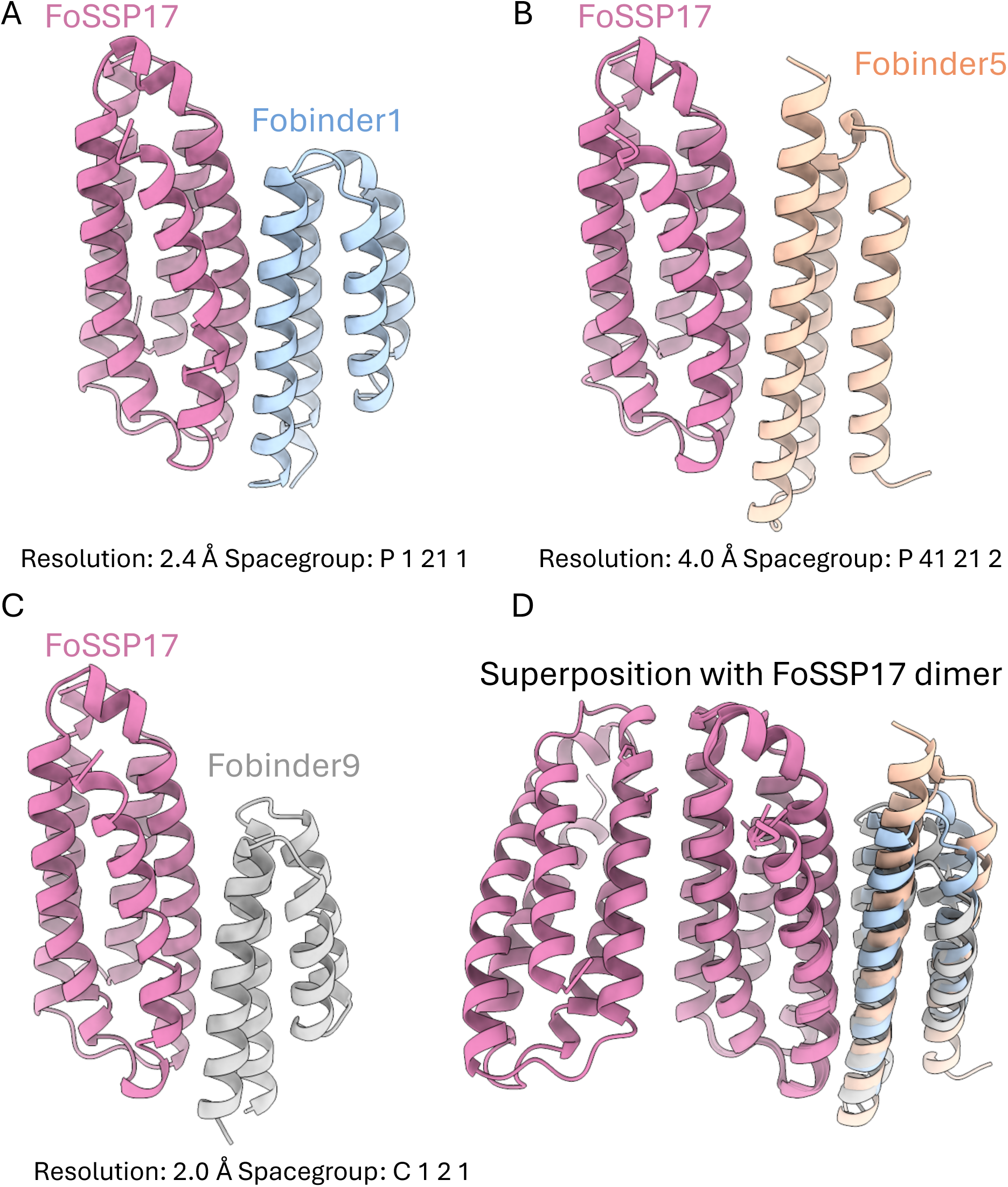
Crystal structures of Fobinder1, 5 and 9 in complex with FoSSP17 demonstrate similar interaction interface. Crystal structures of FoSSP17 in complex with Fobinder1 (A), Fobinder5 (B), and Fobinder9 (C) were obtained at resolutions of 2.40 Å, 4.0 Å, and 2.0 Å, respectively. (D) Superposition of the three complexes with the FoSSP17 homodimer, showing that each Fobinder associates with a similar interface on FoSSP17, mediated by the residues of the α3 and α5 helices of the effector.

**Figure S11.**
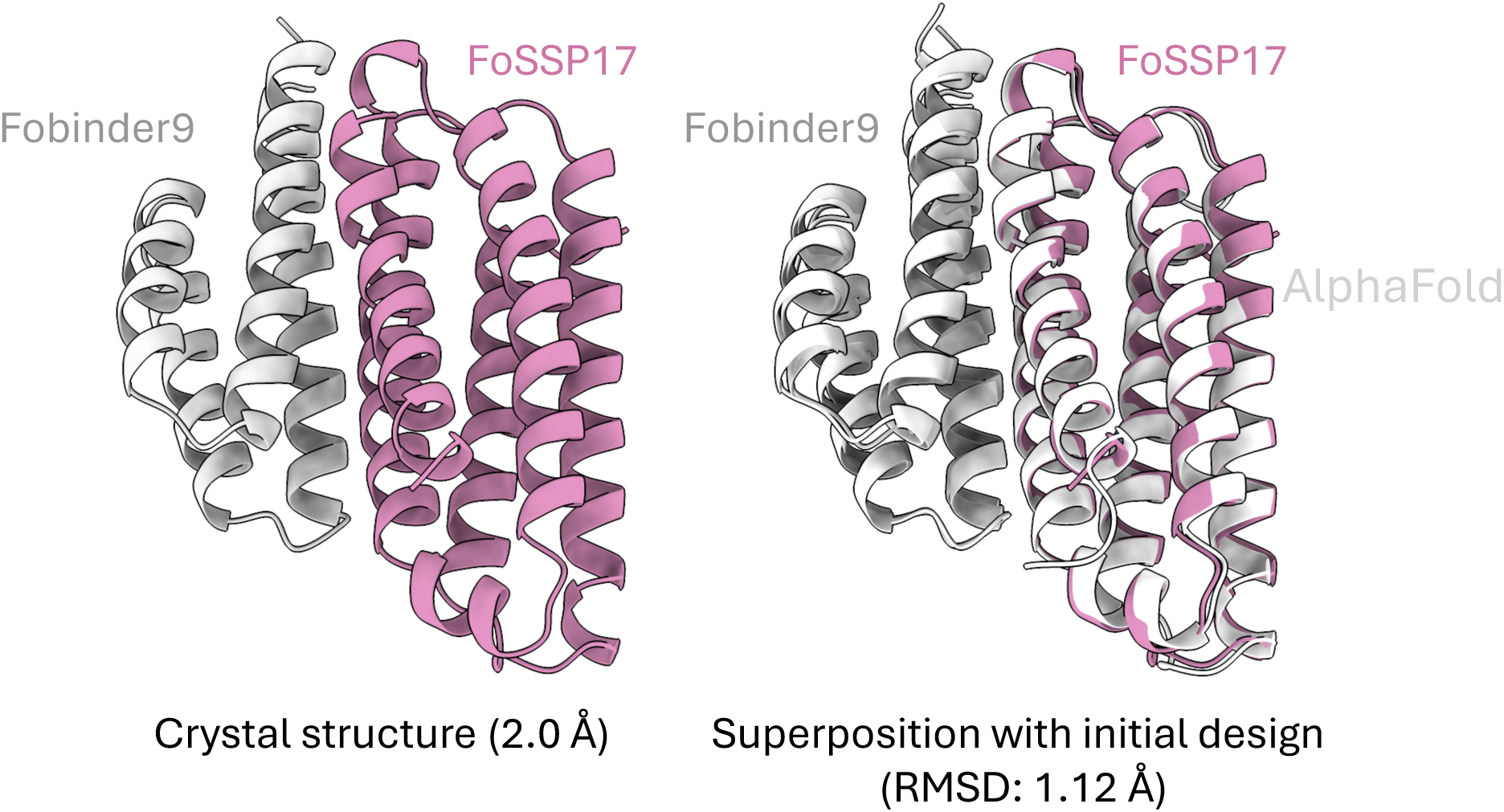
The crystal structure of the Fobinder9/FoSSP17 complex shows high fidelity to initial design model. The crystal structure of the Fobinder9/FoSSP17 complex is shown on the left and was resolved at 2 Å resolution. A superposition of the crystal structure with the AlphaFold2-predicted model of the initial design is shown on the right, with an RMSD of 1.120 Å, indicating reasonable structural similarity between the predicted and experimentally determined complex.

**Figure S12.**
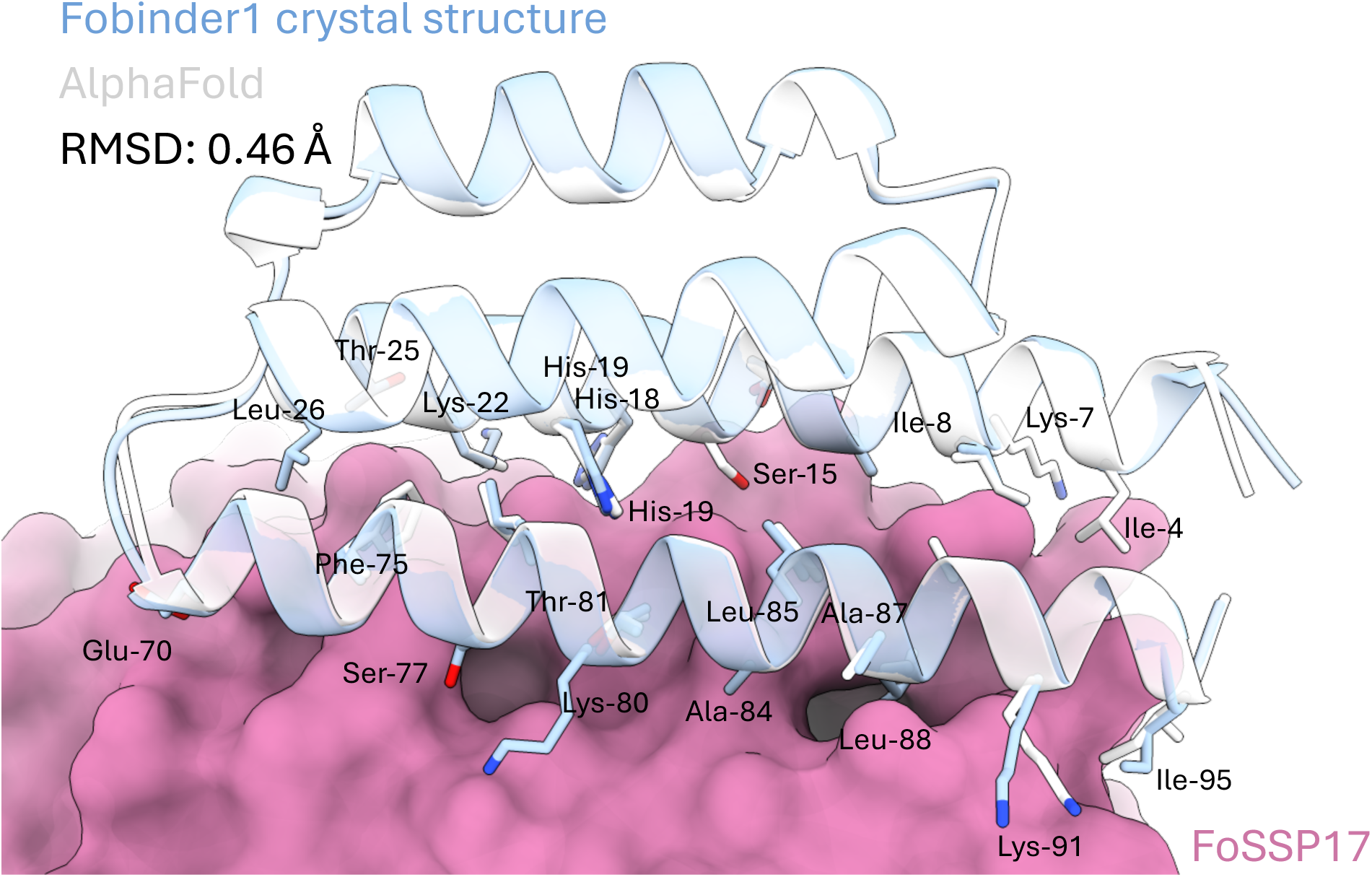
Molecular interactions that govern Fobinder1 interaction with FoSSP17 compared to original binder design with RFdiffusion as predicted with AlphaFold2. Superposition of the AlphaFold2-predicted model of the initial Fobinder1 design with the experimentally determined crystal structure of the Fobinder1/FoSSP17 complex (RMSD = 0.46 Å), shown against the FoSSP17 surface (pink). Side chains of Fobinder1 residues contacting FoSSP17 are shown as sticks and labelled (Ile-4, Lys-7, Ile-8, Ser-15, His-18, His-19, Lys-22, Thr-25, Leu-26, Glu-70, Phe-75, Ser-77, Lys-80, Thr-81, Ala-84, Leu-85, Ala-87, Leu-88, Lys-91, Ile-95), demonstrating close agreement in side-chain placement between the design model and the experimentally determined structure.

**Figure S13.**
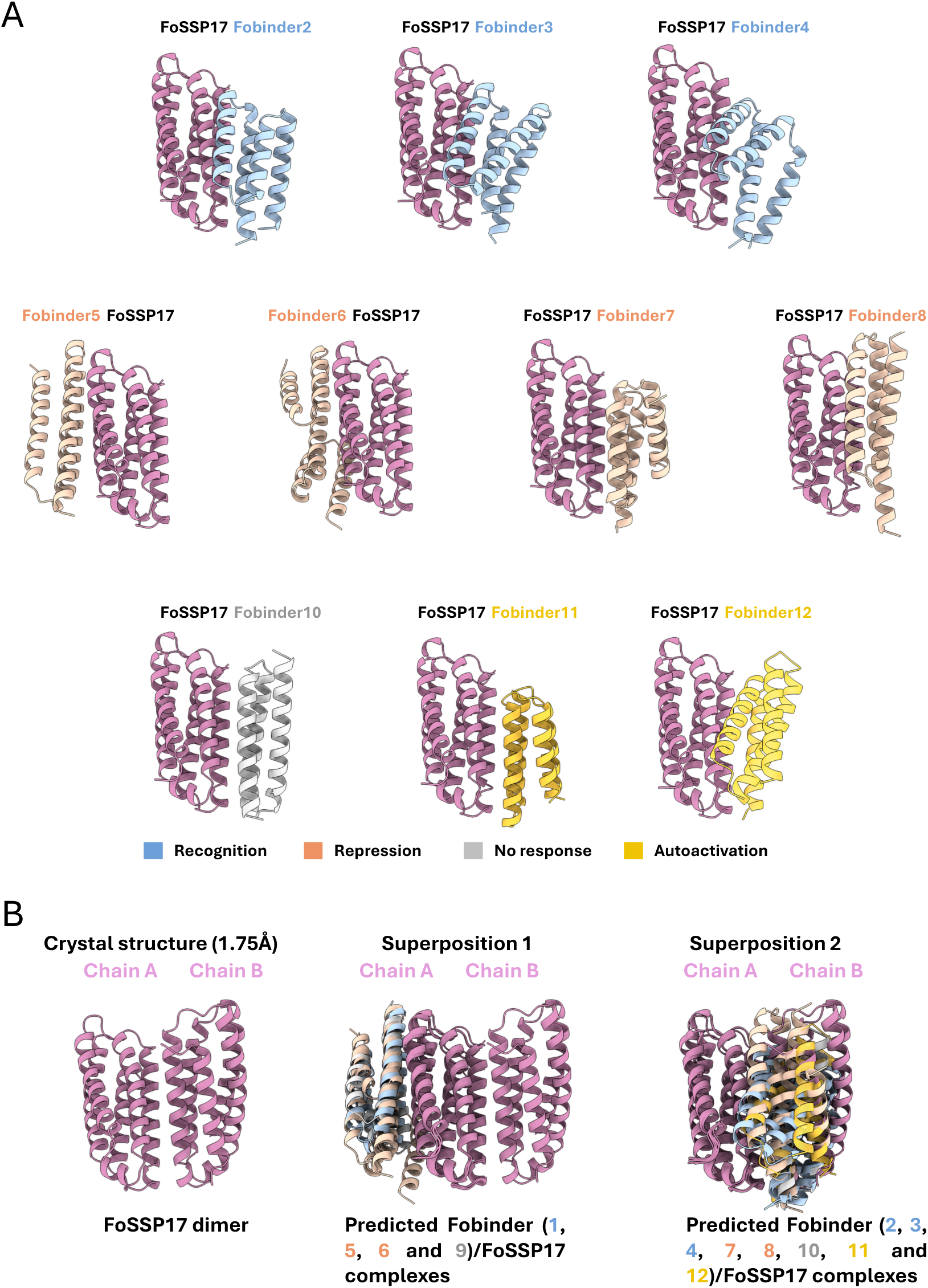
The dimerisation interface of FoSSP17 may occlude some FoSSP17/Fobinder interactions in vitro. (A) AlphaFold2-predicted models of FoSSP17 in complex with each of the ten Fobinders not covered by experimental structures (Fobinder2, 3, 4, 5, 6, 7, 8, 10, 11 and 12; coloured by phenotype: Recognition, blue; Repression, salmon; No response, grey; Autoactivation, yellow). (B) The crystal structure of the FoSSP17 dimer (Chain A, Chain B) is shown on the left and was resolved at 1.75 Å resolution. In the middle, the FoSSP17 dimer is superimposed with the AlphaFold2-predicted Fobinder1, 5, 6 and 9/FoSSP17 complexes, which are predicted to engage the same face of FoSSP17. On the right, the FoSSP17 dimer is superimposed with the AlphaFold2-predicted Fobinder2–4, 7, 8 and 10–12/FoSSP17 complexes, which are predicted to engage the opposing, dimer-occluded face. These superpositions indicate that the binding interfaces of the latter group overlap with the FoSSP17 dimerisation interface in solution, which may explain why only three Fobinders (1, 5 and 9) produced a co-eluting peak in analytical size-exclusion chromatography (Figure S5).

**Table S1.** Summary of Fobinder in planta and biochemical characterization with FoSSP17.

| Binder | Cell death screening (in the Pik chassis) | Co-IP assays | Expression in *E.coli* | Complex in analytical size exclusion chromatography | Spectral shift | Crystal structure |
| --- | --- | --- | --- | --- | --- | --- |
| Fobinder1 | Weak recognition | Association | Stable | Yes | Binding | Solved |
| Fobinder2 | Autoactive but showing recognition | Association | Stable | No | - | - |
| Fobinder3 | Autoactive but showing recognition | Association | Stable | No | - | - |
| Fobinder4 | Strong recognition | Association | Stable | No | Binding | - |
| Fobinder5 | Repression | Association | Stable | Yes | Binding | Crystallised but unable to be refined |
| Fobinder6 | Repression | Association | Unstable | - | - | - |
| Fobinder7 | Repression | Association | Stable | No | - | - |
| Fobinder8 | Repression | Association | Stable | No | - | - |
| Fobinder9 | No response | Association | Stable | Yes | Binding | Solved |
| Fobinder10 | No response | Association | Stable | No | - | - |
| Fobinder11 | Autoactivation | Association | Stable | No | - | - |
| Fobinder12 | Autoactivation | Association | Stable | No | Binding | - |

**Table S2.**
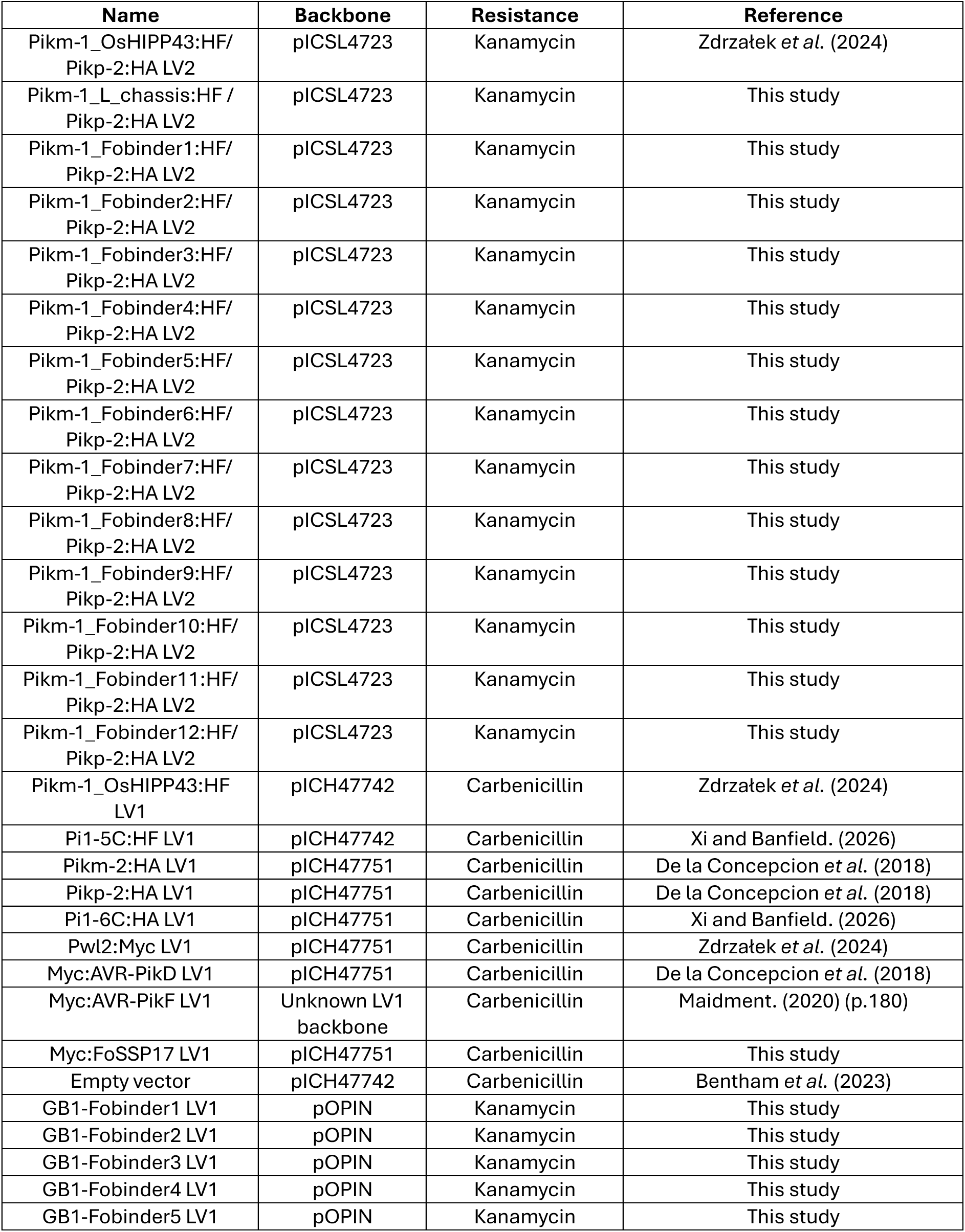
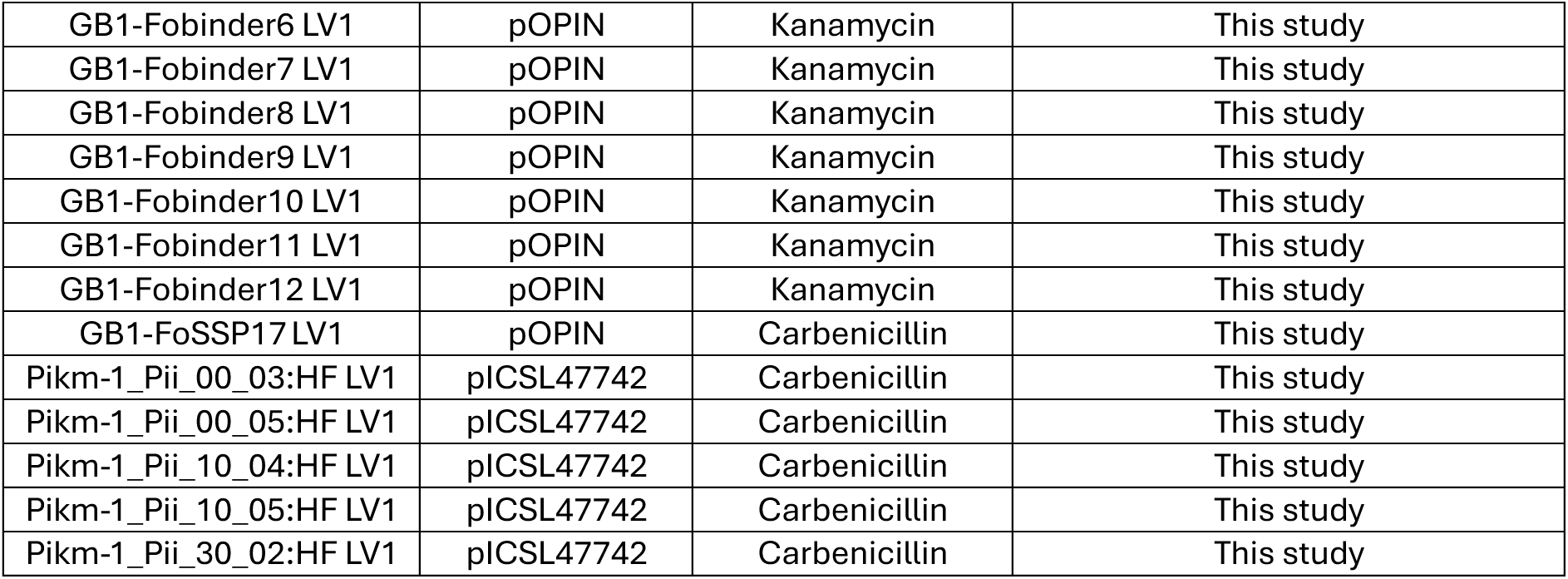
List of constructs used in this study.

**Table S3.**
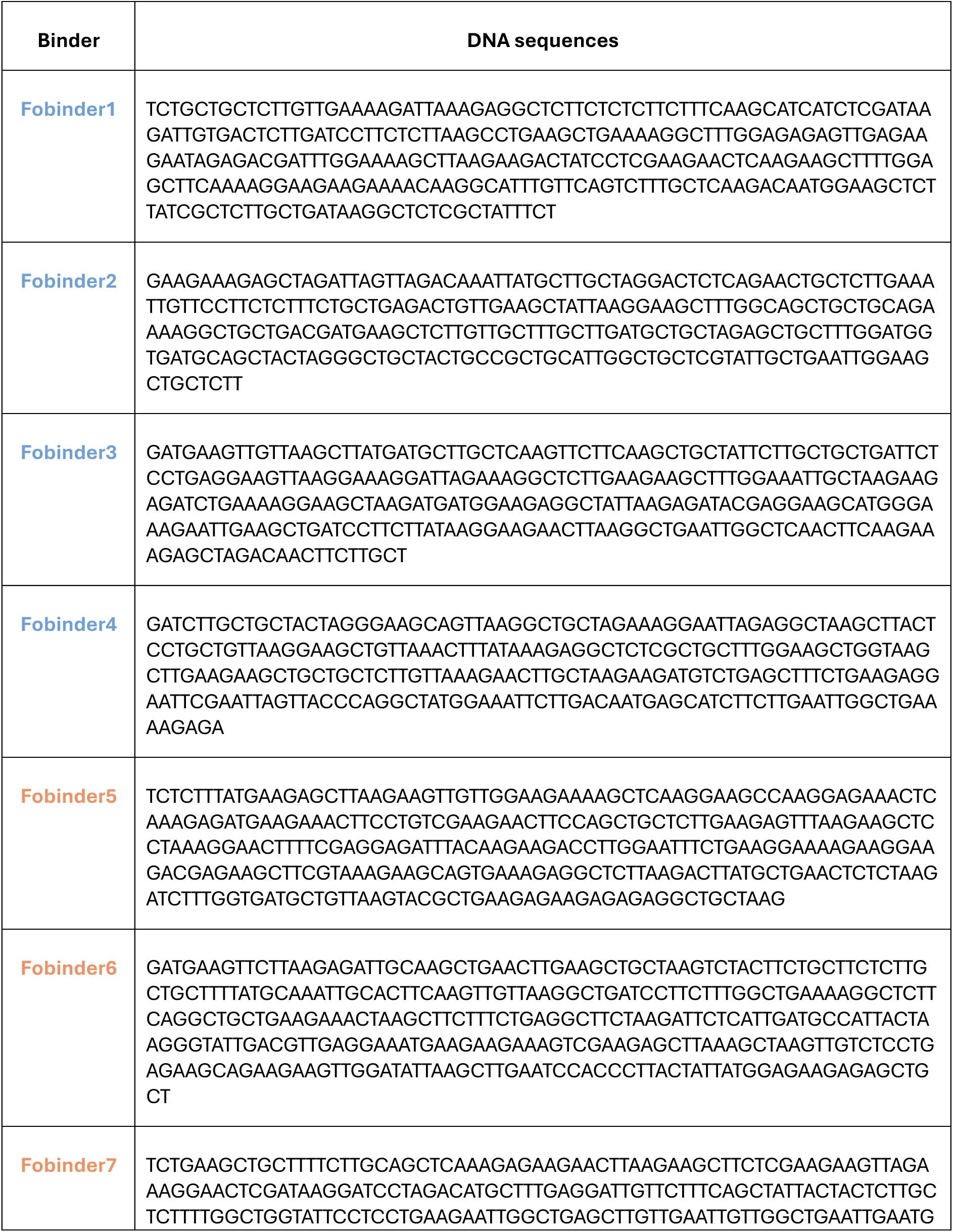
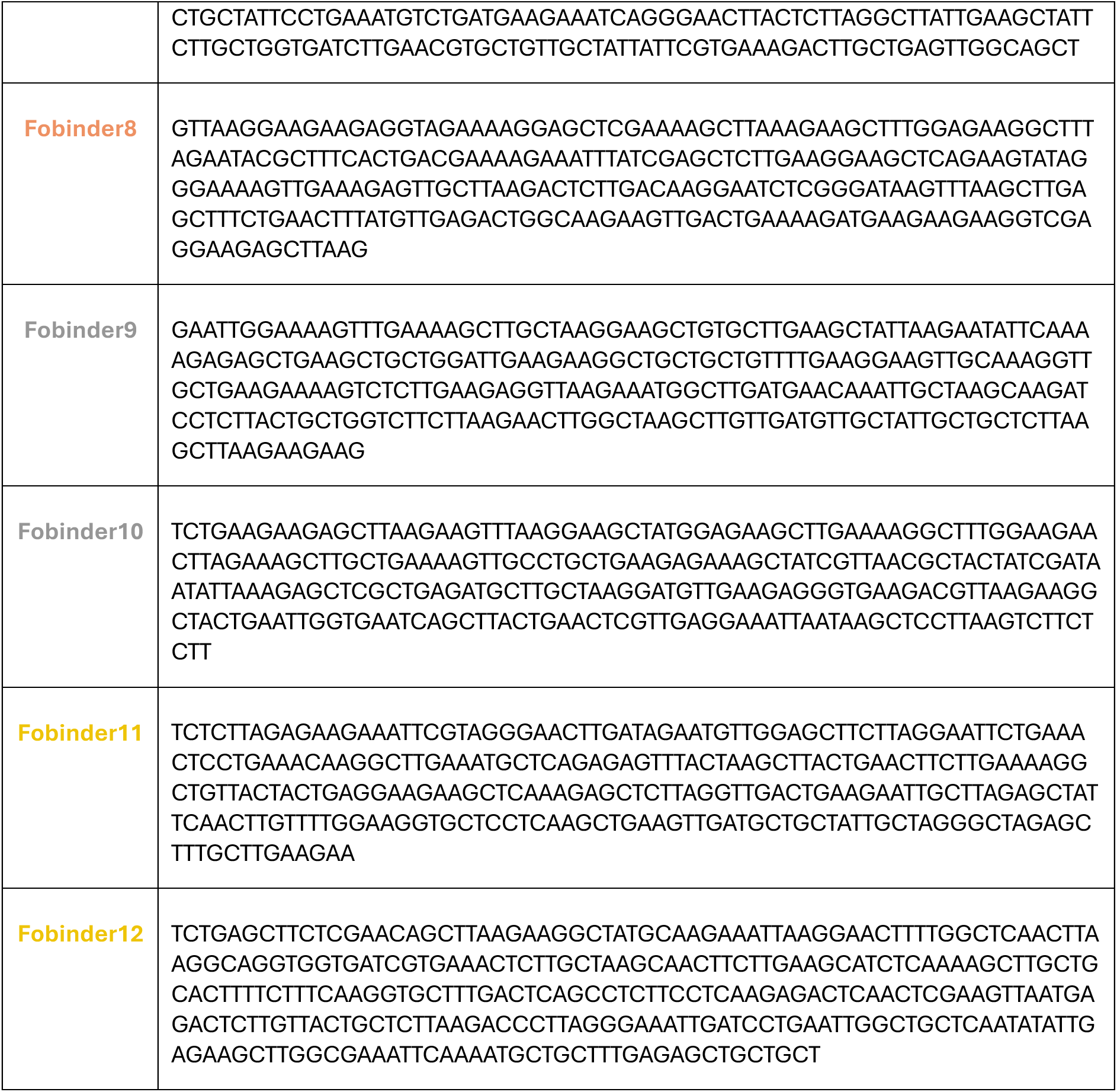
Fobinder sequences generated in this study.

**Table S4.**
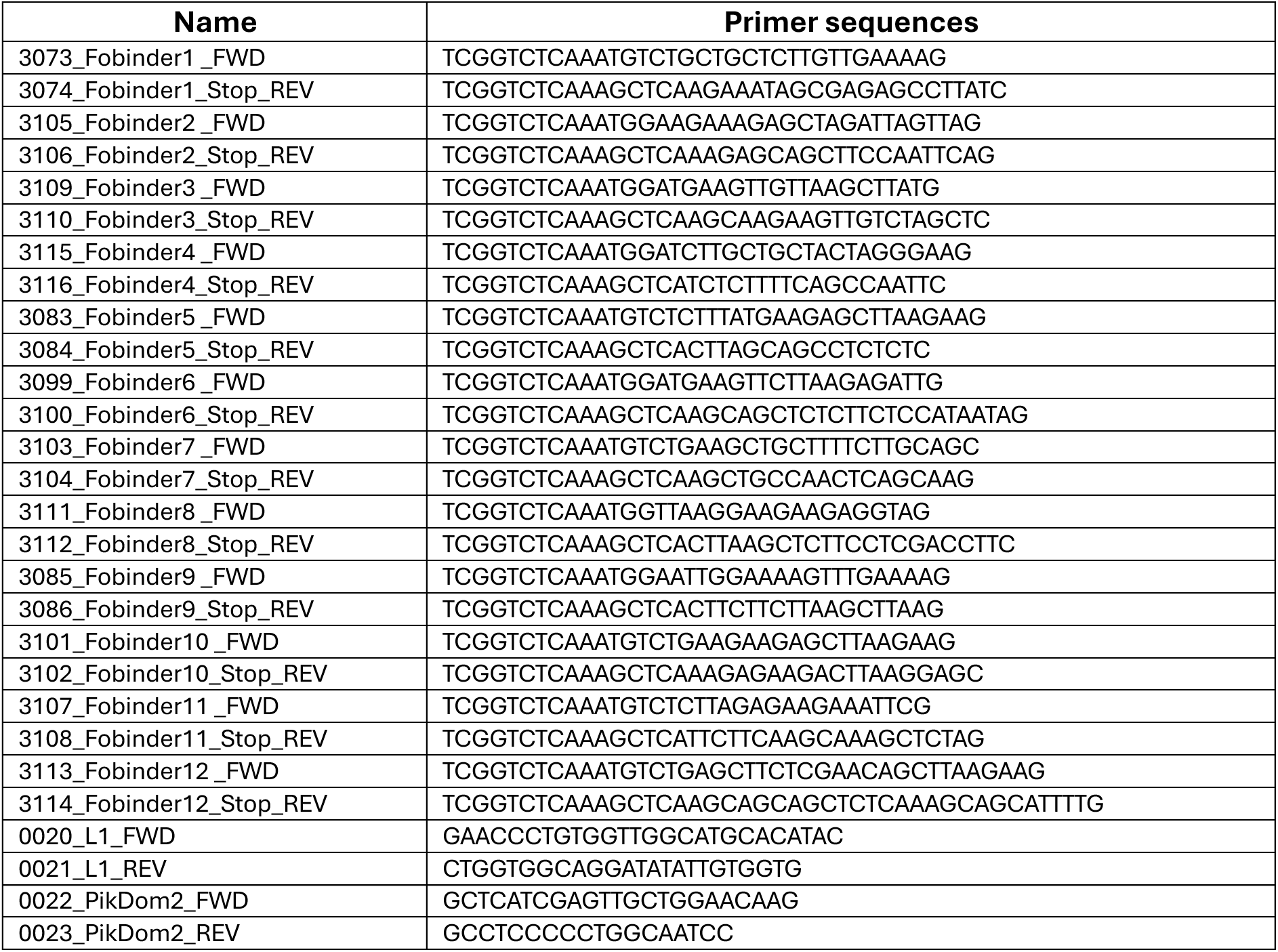
Primers used in this study.

**Table S5.** X-ray data collection and refinement statistics.

|  | FoSSP17/Fobinder1<br>(32EX) | FoSSP17/Fobinder9<br>(32EY) | FoSSP17 dimer<br>(32EW) |
| --- | --- | --- | --- |
| **Data collection statistics** |  |  |  |
| Wavelength (Å) | 0.953730 | 0.953731 | 0.619898 |
| Space Group | P 1 21 1 | C 1 2 1 | P 2 21 21 |
| Unit cell | 41.250 113.670 45.650<br>90.000 95.600 90.000 | 72.449 53.397 99.635<br>90.000 102.839 90.000 | 45.564 67.028 97.210<br>90.000 90.000 90.000 |
| **a, b, c (Å)** |  |  |  |
| Resolution (Å)* | 56.83-2.4 (2.49-2.4) | 48.57-2 (2.05-2) | 67.03-1.75 (1.78-1.75) |
| $R_{meas}$ | 0.135 (1.508) | 0.137 (1.19) | 0.131 (1.627) |
| $I/\sigma I$ | 11.7 (2.2) | 13.1 (2.6) | 14.3 (2) |
| **Completeness (%)** |  |  |  |
| Overall | 100 (100) | 100 (100) | 100 (100) |
| Anomalous | 99.9 (100) | 100 (100) | 100 (100) |
| Unique reflections | 16402 (1688) | 476088 (31570) | 25294 (1880) |
| **Multiplicity** |  |  |  |
| Overall | 14 (14.3) | 18.8 (16.8) | 25.3 (23) |
| Anomalous | 7.1 (7.2) | 9.6 (8.5) | 13.1 (11.7) |
| $CC_{(1/2)}$ | 0.999 (0.864) | 0.999 (0.855) | 0.999 (0.871) |
| **Refinement and model statistics** |  |  |  |
| Resolution (Å) | 45.47-2.4 | 48.62-2.00 | 55.24-1.75 |
| $R_{work}/R_{free}$ | 0.234/0.305 | 0.198/0.257 | 0.184/0.233 |
| **No. atoms** |  |  |  |
| Protein | 7841 | 6172 | 4692 |
| Ligands | - | - | 102 |
| Water | 14 | 119 | 249 |
| **B-factors** |  |  |  |
| All atoms (Å<sup>2</sup>) | 65 | 41 | 34 |
| **R.m.s deviations** |  |  |  |
| Bond lengths (Å) | 0.0057 | 0.012 | 0.0106 |
| Bond angles (°) | 1.576 | 2.181 | 1.909 |
| **Ramachandran Plot\*\*** |  |  |  |
| Favoured | 466 | 380 | 295 |
| Allowed | 17 | 5 | 2 |
| Outliers | 2 | 2 | 0 |
| **MolProbity Score** | 2.12 | 1.75 | 1.22 |

\*The highest resolution shell is shown in parenthesis
\*\*As calculated by MolProbity

## Resource Availability

### Lead Contacts

Further information and requests for resources and reagents should be directed to and will be fulfilled by the lead contacts, Adam Bentham and Mark Banfield.

### Materials Availability

All constructs generated in this study will be made available upon request, pending completion of a Materials Transfer Agreement (MTA).

### Data and Code Availability

Atomic coordinates and structure factors for the crystal structures reported in this paper have been deposited in the Protein Data Bank under accession codes 32EW, 32EX, and 32EY. This paper does not report original code.

Any additional information required to reanalyze the data reported in this paper is available from the lead contact upon request.

## Acknowledgments

We thank the staff at Diamond Light Source (Oxford, UK) for access to and support with X-ray data collection facilities and thank the members of the John Innes Centre Structural Biology Platform, Prof. David Lawson, Julie Mundy and Dr. Sandra Eltschkner for their assistance in data collection. Thank you to Dr. Mark Youles of TSL SynBio in his assistance with the generation several constructs used in this study. This work was supported by the Rank Prize Funds to ARB, grants from UKRI-BBSRC (BB/X010996/1, BB/V015508/1, BB/W00108X/1 UKRI1913), the BBSRC Norwich Research Park Biosciences Doctoral Training Partnership (grant BB/T008717/1), and John Innes Foundation (PhD Rotation Programme) to MJB, a Leverhulme International Professorship (grant LIP-2022-010) awarded by the Leverhulme Trust to JGH, and grants from the Gatsby Charitable Foundation and UKRI-BBSRC (BB/X010996/1, BB/Y002997/1) to NJT.

## Author Contributions

Conceptualization, YX, AHB, PE, JLW, NJT, MJB, and ARB.; Methodology, YX, AHB, MJB, and ARB.; Investigation, YX, AHB, JLW, AM, JWB, IG, DV, GK, RZ, CR, IS, CES, EKT, DSY, AG, LSR, XY, VW, and ARB; Formal Analysis, YX, AHB, JLW, AM, MJB, and ARB; Writing – Original Draft, YX and ARB.; Writing – Review & Editing, all authors; Visualization, YX, AHB, and ARB.; Supervision, DB, NJT, MJB, and ARB.; Funding Acquisition, ARB, JGH, NJT, and MJB.

## Declaration of Interests

The authors declare no competing interests.

## Materials and Methods

### Binder design

Protein binder design campaigns were performed with RFdiffusion⁷ against three effector targets: AVR-PikF, AVR-Pii and FoSSP17. Existing experimental structures were used as design targets for AVR-PikF²⁹ and AVR-Pii³⁰. As no experimental structure existed for FoSSP17 at the time of this study, a high-confidence AlphaFold2³¹ v1.5.5 monomer prediction, generated as implemented in ColabFold³² (pLDDT > 0.90; pTM > 0.85), was used as the design target input (Figure S1).

For designing binders against AVR-PikF and FoSSP17, we used a slight modification of the binder design protocol described in Watson et al., 2023^7^, with the goal of maximising the diversity of designs generated. 19200 initial designs were generated for each target, representing a grid search over noise scales (0, 0.5, 1.0), fold-conditioned (50% of designs unconditional, 50% conditioned on pre-defined scaffolds), and checkpoints (four RFdiffusion checkpoints were used from different stages of training). For further diversity and cost-efficiency, the final 10 timepoints of the denoising trajectory were extracted. Following backbone generation, ProteinMPNN^6^ was used to design a single low-temperature sequence (T=0.0001) on each of these 10 backbones. AlphaFold2^33^ with initial guess^52^ was used to subsequently re-predict the structure. The total number of designed sequences per target was therefore 192,000. Of these, the top 5000 by pAE (after filtering by R.M.S.D. to design structure < 2Å) were further refined by partial diffusion^9^ with three different renoising parameters (15, 20, 25, out of a possible 50 in the default RFdiffusion schedule), yielding approximately 100,000 optimised variants. From these, sequences were clustered at appropriate sequence similarity using MMSeqs2, and the top approximately 10,000 were selected for subsequent validation by yeast surface display as per the methods described in Bennet et al. 2024^5^.

For AVR-Pii, 256 backbone structures were generated with RFdiffusion v1.1.1 using the ColabDesign implementation^35^ (github.com/sokrypton/ColabDesign). Each backbone was used as a template for inverse folding with ProteinMPNN using a sampling temperature of 0.0001 and with cysteines excluded from the design, providing eight sequences per backbone and resulting in a total of 2048 binder designs. Binder and effector sequences were subjected to structure prediction with AlphaFold2 in single-sequence mode (no multiple sequence alignment), with the target protein provided via the template feature and the RFdiffusion design model used as an initial guess, following established best practice^7^. Designs were filtered based on predicted confidence metrics (pLDDT > 0.85; ipTM > 0.80; ipAE < 5.0) to select candidates for experimental validation.

### Constructs

For agroinfiltration assays, we generated a new LV2 2×CaMV35S:Pikm-1_RFP cassette:HF/mas:Pikp-2:HA construct for Golden Gate (GG) cloning, following a strategy similar to that described previously^27^. Differently, in the new 2×CaMV35S:Pikm-1_RFP cassette:HF module, partial N– and C-terminal sequences of the wild-type Pikm-1^HMA^ region were retained rather than replacing the entire Pikm-1^HMA^ domain with the RFP selection cassette (Table S2), with the aim of reducing the emergence of Pik pair autoactivity after engineering. FoSSP17 (signal peptide excluded, from amino acid 18 to 172) and all designed Fobinders (the DNA sequences of selected Fobinders are listed in Table S3), carrying compatible BsaI overhangs, were synthesised as gBlock gene fragments by Integrated DNA Technologies (IDT) and cloned into the corresponding backbones for cell death assays via GG assembly (Table S2). Apart from Pwl2, which carried a C-terminal Myc tag, all other effectors were fused with an N-terminal Myc tag. All effectors used in this study were driven by the *AtUbi10* promoter.

For in vitro protein expression of the selected Fobinders, primers were designed (Table S4) and synthesised gene fragments were used as templates for PCR amplification. The purified PCR products were subsequently assembled into the pOPIN backbone with an N-terminal His-GB1 tag via GG assembly. GB1-FoSSP17 (carrying His-GB1 tag) was generated by directly cloning the synthesised gene fragment described above into the pOPIN backbone via GG assembly (Table S2). A 3C protease cleavage site was placed between the GB1 tag and the protein of interest for subsequent tag cleavage. All constructs were verified by Sanger sequencing prior to use in experiments.

### Nicotiana benthamiana cell-death assays

The corresponding constructs prepared for the agroinfiltration assays were all transformed into *Agrobacterium tumefaciens* GV3101 and four-week-old *N. benthamiana* plants were used for cell death assays. GV3101 agrobacteria carrying LV2 constructs and LV1 constructs were cultured at 28°C in LB medium containing kanamycin/rifampicin/gentamicin and carbenicillin/rifampicin/gentamicin, respectively. The infiltration buffer containing 10 mM MgCl₂, 10 mM MES (pH 5.6) and 150 µM acetosyringone was used for agrobacteria resuspension. For the cell death of engineered Pik pair LV2 constructs, LV2 constructs, effectors/empty vector and the P19 were mixed at OD₆₀₀ of 0.5, 0.5 and 0.1, respectively. For the cell death assays of Pi1-5C/Pi1-6C, NLR, effectors/empty vector and the P19 were also mixed at OD₆₀₀ of 0.5, 0.5 and 0.1, respectively, but when mixing the Pi1-5C and Pi1-6C, both of them were diluted to OD₆₀₀ of 0.25 to balance the total concentration. Leaves were harvested and imaged under both normal and UV light at five days post infiltration, with camera settings as detailed in Bentham *et al.* (2023)^3^.

The cell death symptoms were scored from index 0 to 6 according to the scale as described in the supplementary of De la Concepcion *et al.* (2018)^17^. The cell death score data of Figure 2 C and Figure 4 C are shown in Supplemental Dataset 1. The dot plot analyses were plotted with R v. 4.3.0 (https://www.r-project.org) using the ggplot2 package^53^.

### Protein extraction from plant tissues and western blot

Agrobacterium strains carrying each construct used in this study were individually infiltrated into *N. benthamiana* leaves for protein expression analysis, using OD_600_=0.5 for most constructs, with p19 additionally included at OD_600_=0.1. Since Myc:FoSSP17 and Myc:AVR-PikD showed much stronger expression than Pwl2:Myc in our preliminary tests, both constructs were diluted to OD_600_=0.1. As infiltration with agrobacteria carrying some LV2 constructs resulted in cell death symptoms at a very early stage, five leaf discs were harvested into a 1.5 ml Eppendorf tube at 40 hours post-infiltration and immediately frozen in liquid nitrogen. As previously described^20^, leaf tissues were ground and the powder was resuspended in 200 μl extraction buffer (25 mM Tris-HCl (pH 7.5), 1 mM ethylenediaminetetraacetic acid (EDTA), 150 mM NaCl, 0.1% NP-40 (Sigma), 10% glycerol, 10 mM dithiothreitol (DTT), 0.5% (w/v) polyvinylpolypyrrolidone (PVPP), and 1x protease inhibitor cocktail (Sigma)). The supernatants were then collected by centrifugation at 14000g for 30 minutes at 4°C to obtain protein extracts.

For SDS-PAGE, 8 μl of protein extract was mixed with 8 μl of 2x protein loading dye and loaded onto the gel. Following electrophoresis, proteins were transferred onto a PVDF (polyvinylidene difluoride) membrane using a Trans-Blot Turbo transfer system (Bio-Rad). Prior to western blotting, fresh TBST buffer (50 mM Tris-HCl (pH 8.0), 150 mM NaCl, and 0.1% Tween-20) was prepared. Membranes were blocked with 5% (w/v) skimmed milk in TBST for 1 hour after protein transfer, followed by incubation with HRP-conjugated primary antibodies (α-FLAG, Cohesion Biosciences; α-HA, Invitrogen; Rabbit α-Myc, Bethyl) diluted 1:3000 for a further 1 hour. To remove non-specifically bound antibody, membranes were washed four times for 5 minutes each with TBST, and signals were detected using an ImageQuant LAS 500 imager (GE Healthcare) after adding Clarity Max Western ECL Substrate (Bio-Rad). Total protein loading was monitored by Ponceau staining.

### In planta co-immunoprecipitation assays

Agrobacterium strains carrying LV2 constructs and target effectors were transiently co-expressed in *N. benthamiana*, using OD_600_=0.5 for both LV2 constructs and effectors, and OD_600_=0.1 for p19. As co-expression of Pikm-1_HIPP43:HF/Pikp-2:HA with Pwl2:Myc induced strong cell death at 24 hours post-infiltration, we instead co-expressed the Pikm-1_HIPP43:HF LV1 construct with effectors as controls. At 40 hours post-infiltration, five leaf discs were harvested into Eppendorf tubes and immediately frozen in liquid nitrogen. Frozen leaf samples were ground, and the powder was resuspended in 600 μl IP extraction buffer (the same buffer used in the “protein extraction from plant tissues” section, with Roche protease inhibitor tablets added at 1 tablet/50 ml). The supernatants were then collected by centrifugation at 14000g for 30 minutes at 4°C. To prepare input samples, 10 μl of each extract was mixed with 10 μl of 2x protein loading dye and denatured at 95°C for 10 minutes. The remaining protein extracts were mixed with 6 μl of anti-FLAG M2 magnetic beads (Sigma, M8823) and incubated for at least 1 hour on a rotary mixer at 4°C. To remove non-specifically bound proteins, the anti-FLAG magnetic beads were washed four times with washing buffer (IP extraction buffer without adding PVPP, DTT and protease inhibitor cocktail (Sigma)). To prepare IP samples, the washed beads were incubated with 50 μl of 1x protein loading dye at 70°C for 10 minutes, and the supernatants were collected after separation on a magnetic rack. 20 μl input samples and 12 μl IP samples were separated by SDS-PAGE and analysed by western blot.

### Protein expression and purification from *E. coli*

The relevant constructs were all transformed into SHuffle T7 Express competent cells, and the transformants were grown on LB agar plates supplemented with the appropriate antibiotics at 30°C (GB1-Fobinders: kanamycin; GB1-FoSSP17: carbenicillin). To induce protein production, selected colonies were cultured in LB liquid medium with the appropriate antibiotics and incubated in a shaker at 30°C and 220 rpm until the OD_600_ reached 0.6-0.8. Isopropyl β-D-1-thiogalactopyranoside (IPTG) was then added to a final concentration of 1 mM, and the incubation conditions were shifted to 16°C and 180 rpm for overnight growth (no more than 20 hours). Cell pellets were harvested by centrifugation at 5500g for 10 minutes at 4°C and resuspended in lysis buffer containing 50 mM HEPES (pH 8.0), 50 mM glycine, 500 mM NaCl, 5% (v/v) glycerol and 20 mM imidazole, freshly supplemented with cOmplete EDTA-free Protease Inhibitor Cocktail (1 tablet/ 50 ml).

The resuspended cells were then sonicated (40% amplitude, 1 s on/3 s off) until homogeneous, and the supernatants were collected by centrifugation at 43000g for 30 minutes at 4°C for downstream purification. To purify the single His-GB1-tagged proteins, the supernatants were loaded onto a 5 ml HisTrap FF nickel column (Cytiva), washed with lysis buffer to remove unbound proteins (10 column volumes), and eluted in a single step with high-imidazole buffer (50 mM HEPES (pH 8.0), 50 mM glycine, 500 mM NaCl, 5% (v/v) glycerol and 500 mM imidazole). The eluted proteins were then buffer-exchanged into a buffer containing 20 mM HEPES (pH 7.5) and 150 mM NaCl using a SepFast 26/10 Desalting Column (BioToolomics Ltd, UK). To remove the His-GB1 tags, the protein solutions were incubated with 3C protease overnight at 4°C, and the cleaved His-GB1 tag was separated from the proteins of interest by reloading the mixture onto the 5 ml nickel column and collecting the flowthrough. Proteins were further purified by size-exclusion chromatography (SEC) with the same buffer used for desalting. The proteins of interest in each step were confirmed by SDS-PAGE, and the untagged proteins were collected and concentrated before storage at −70°C for future use.

### Analytical size-exclusion chromatography

Purified FoSSP17, Fobinders (Fobinder6 was excluded due to its instability) and FoSSP17/Fobinder mixtures were loaded onto a Superdex 75 Increase 5/150 GL (Cytiva) column at 4°C for separation, and UV absorbance at 280 nm was monitored. The FoSSP17/Fobinder mixtures were incubated on ice for at least 1 hour before loading. A buffer containing 20 mM HEPES (pH 7.5) and 150 mM NaCl was used for column equilibration. Eluted fractions were collected for SDS-PAGE analysis. Of note, Fobinder1, 2, 7, 10 and 11showed no UV absorbance at 280nm.

### Crystallisation, x-ray data collection and structure determination

To prepare Fobinder1/FoSSP17, Fobinder5/FoSSP17 and Fobinder9/FoSSP17 complexes for crystallisation screening, purified FoSSP17 and the corresponding Fobinders were mixed and incubated on ice for at least 1 hour. The mixtures were then rerun by SEC and the peak fractions were collected for crystallisation trials. Screening was performed by sitting-drop vapour diffusion (0.3 µl protein solution plus 0.3 µl reservoir solution) in 96-well plates using an Oryx nano robot (Douglas Instruments, UK), and sealed plates were incubated at 18°C. Crystals of FoSSP17 dimer, Fobinder1/FoSSP17 and Fobinder9/FoSSP17 complexes were obtained at protein concentrations of 11, 7 and 9.5 mg/ml, respectively, all from the Morpheus kit screen (Molecular Dimensions). Crystals of Fobinder5/FoSSP17 complexes were obtained at protein concentrations of 8 mg/ml from the PROPLEX screen (Molecular Dimensions). Specifically, FoSSP17 dimer crystallised in 0.1 M Carboxylic acids, 0.1 M Buffer System 3 (pH 8.5) and 50 % (v/v) Precipitant Mix 1; Fobinder1/FoSSP17 crystallised in 0.06 M Divalents, 0.1 M Buffer System 1 (pH 6.5) and 50% (v/v) Precipitant Mix 2; Fobinder5/FoSSP17 in 0.1 M Imidazole pH 7.0 and 50 % (v/v) MPD 1; and Fobinder9/FoSSP17 in 0.06 M Divalents, 0.1 M Buffer System 3 (pH 8.5) and 50% (v/v) Precipitant Mix 4. Crystals were harvested and briefly soaked in the respective crystallisation condition supplemented with 20% (v/v) ethylene glycol as cryoprotectant (except for FoSSP17, which required no additional cryoprotectant), and snap cooled in liquid nitrogen.

X-ray diffraction data were collected on the I04 and I24 beamlines at the Diamond Light Source (Oxford, UK). Data were scaled and merged using the AIMLESS^54^ pipeline in CCP4i2^55,56^. All structures were solved by molecular replacement using PHASER^57^. The FoSSP17 dimer structure was solved using AlphaFold2^33^-predicted models (v1.5.5) as a template model, and this structure was subsequently used as a template for solving the Fobinder/FoSSP17 complex structures, alongside AlphaFold2-predicted models of Fobinder1, 5 and 9 (v1.6.1). Iterative rounds of manual model building in COOT^58^ and refinement with REFMAC^59^ as implemented in the CCP4i2 crystallographic software suite^55,56^, were carried out until satisfactory geometry and R-factors were achieved. X-ray data collection and refinement statistics are shown in Table S5. Final structures for the FoSSP17 dimer, Fobinder1/FoSSP17 complex and Fobinder9/FoSSP17 complex have been deposited in the Protein Data Bank (PDB) under accession codes 32EW, 32EX, and 32EY, respectively.

### Spectral shift assays

Binding affinities between Fobinders and FoSSP17 were determined by spectral shift assay using Monolith X instrument (NanoTemper Technologies). As labelled FoSSP17 could not be eluted from the column following conjugation, the five Fobinders used in this study were labelled individually instead. Fobinders were labelled with the Protein Labelling Kit RED-NHS 2nd Generation (NanoTemper Technologies, Cat# MO-L011) according to the manufacturer’s instructions. Labelled Fobinders were used at a final concentration of 20-30 nM, and FoSSP17 was titrated as the ligand at concentrations ranging from 50 µM to 850 µM across a 16-point dilution series. All experiments were performed in triplicate in assay buffer containing 20 mM HEPES (pH 7.5), 150 mM NaCl and 0.05% Tween-20. Fluorescence emission was monitored at 650 nm and 670 nm, and the ratio of emission intensities (670/650 nm) was plotted against ligand concentration. Dissociation constant (*K_d_*) values were determined by fitting each replicate independently to a 1:1 binding model, and the mean value with standard deviation is reported.

### Right-angle laser light scattering

Right-angle laser light scattering (RALS) coupled with refractive index (RI) detection was performed using a Malvern Panalytical OMNISEC system. 20 µl of FoSSP17 at 500 µM was separated on a Superose 6 10/300 Increase analytical size-exclusion column (Cytiva) in 20 mM HEPES pH 7.4, 150 mM NaCl, 1 mM DTT, at a flow rate of 0.5 ml/min at 25°C. The light scattering and RI detectors were normalized using bovine serum albumin (BSA) as a calibration standard prior to sample analysis. Light scattering and RI signals across the elution peak were recorded, and molecular mass was calculated using a dn/dc of 0.187 ml/g (per manufacturer’s instructions for unmodified proteins) in the OMNISEC software, assuming a second virial coefficient (A2) of zero. Measurements were performed in three replicate injections.

## Notes

### Competing Interest Statement

The authors have declared no competing interest.

### Summary of Updates

Fixed typos and errors in table numbering.

